# Bicyclic pyrrolidine inhibitors of *Toxoplasma gondii* phenylalanine t-RNA synthetase with antiparasitic potency *in vitro* and brain exposure

**DOI:** 10.1101/2024.02.28.582607

**Authors:** Chloe C. Ence, Taher Uddin, Julien Borrel, Payal Mittal, Han Xie, Jochen Zoller, Amit Sharma, Eamon Comer, Stuart L. Schreiber, Bruno Melillo, L. David Sibley, Arnab K. Chatterjee

**Author notes:** Corresponding Authors: Bruno Melillo,; L. David Sibley,; Arnab Chatterjee.

## Abstract

Previous studies have shown that bicyclic azetidines are potent and selective inhibitors of apicomplexan phenylalanine tRNA synthetase (PheRS), leading to parasite growth inhibition *in vitro* and *in vivo*, including in models of *Toxoplasma* infection. Despite these useful properties, additional optimization is required for the development of efficacious treatments of toxoplasmosis from this inhibitor series, in particular to achieve sufficient exposure in the brain. Here, we describe a series of PheRS inhibitors built on a new bicyclic pyrrolidine core scaffold designed to retain the exit-vector geometry of the isomeric bicyclic azetidine core scaffold while offering avenues to sample diverse chemical space. Relative to the parent series, bicyclic pyrrolidines retain reasonable potency and target selectivity for parasite PheRS vs. host. Further structure-activity relationship studies revealed that the introduction of aliphatic groups improved potency, ADME and PK properties, including brain exposure. The identification of this new scaffold provides potential opportunities to extend the analog series to further improve selectivity and potency and ultimately deliver a novel, efficacious treatment of toxoplasmosis.

**Lay abstract:** The inhibition of protein synthesis in parasites has emerged as an attractive strategy to target parasitic diseases such as malaria, cryptosporidiosis, and toxoplasmosis. In this study, we report a new series of small molecules that inhibit the enzyme responsible for loading the amino acid phenylalanine onto its cognate tRNA in the parasite species *Toxoplasma*, directly upstream of protein synthesis. We show that small molecules in this new series inhibit *Toxoplasma* parasite growth at low concentrations, and that, orally administered to mice, these molecules achieve high concentrations in the brain, which hold promise for the treatment of forms of toxoplasmosis that result from parasitic brain infection.

---

*Toxoplasma gondii* is a common intracellular parasite of companion, agricultural, and wild animals that also infects up to one-third of the world’s human population.^1^ Following initial expansion and dissemination of actively replicating tachyzoites, a vigorous immune response controls parasite replication.^2^ In response to stress, the parasite differentiates into a slow growing, semi dormant stage called the bradyzoite, which resides in cysts in the brain and other tissue.^3^ Such chronic infections pose a risk when deteriorating immune responses fails to control chronic infection and semi-dormant bradyzoites revert to actively replicating tachyzoites, which destroy tissue and cause severe manifestations including encephalitis and vision loss.^4^ Current standard of care treatment of toxoplasmosis targets the folate pathway using a combination of pyrimethamine with sulfadiazine or other sulfa analogs.^5^ Although these formulations are effective at repressing tachyzoite growth, they are inactive on the semi-dormant bradyzoites in the brain and other tissues.^5^ Hence, new treatments are needed to control reactivation and cure chronic infection. To be effective, such medicines would also need to cross the blood brain barrier and not result in complications within this compartment.

We recently found that potent small-molecule inhibitors^6^ of *Plasmodium* cytosolic phenylalanyl tRNA synthetase (cPheRS),^7, 8^ typified by the bicyclic azetidine BRD7929 (Figure 1), also potently inhibit the closely related *Toxoplasma gondii* and *Cryptosporidium* orthologs resulting in antiparasitic activity *in vitro* and efficacy *in vivo*.^9, 10^ This antiparasitic mechanism of action,previously unknown and yet to see clinical introduction in any indication, has garnered attention in the malaria field.^11^ Crystallographic structures of bicyclic azetidines in complex with *Plasmodium falciparum* and *Plasmodium vivax* cPheRS demonstrated that two molecular appendages interacted with deep pockets in the enzyme target: the biarylacetylene motif was found to occupy the L-Phe site, while the methoxyphenyl urea motif occupied a distinct site of unknown function, so-called auxiliary site.^12, 13^ As part of structure-activity relationship (SAR) studies, we envisioned that alternative core scaffolds could mimic the critical exit vectors of the bicyclic azetidine lead series while providing an opportunity to expand the cPheRS inhibitor chemical space. Relying on molecular modeling, we specifically hypothesized that the bicyclic pyrrolidine scaffold in BRD2108 (Figure 1) could position the key appendages in close proximity to those of its bicyclic azetidine isomer BRD3914. The modular synthesis^14^ of BRD3914 was subsequently adapted to access BRD2108 in short order (vide infra Scheme 1).

**Figure 1.**
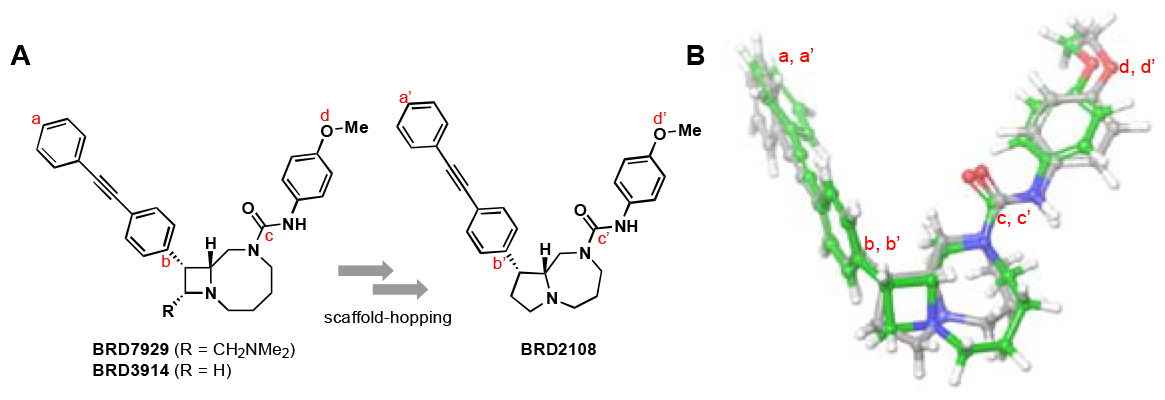
Core-scaffold isomerism provides an opportunity for expansion of cPheRS inhibitor chemical space. **A**. Chemical structure of *Pf* cPheRS inhibitors BRD7929 and BRD3914 (refs. 6, 14) and proposed designed of an isomeric analog, BRD2108. **B**. Superimposition of calculated molecular structures of BRD3914 (green) and BRD2108 (grey). The atoms indicated with red labels were used to quantify the similarity of the positions of key molecular appendages (atom-pair RMSD 0.5813 Å). Ligand geometry optimized using DFT (B3LYP-D3 theory level, 6-31G** basis set, PBF solvation [water]) on Jaguar Version 12.1, Schrödinger Maestro Version 13.7.125, MMshare Version 6.3.125, Release 2023-3, Platform Darwin-x86_64.

We were encouraged to find that BRD2108 inhibited *T. gondii* growth at submicromolar concentrations (EC_50_ 0.27 μM, Table 1), with a significant loss in potency observed in the TgcPheRS L497I mutant, corroborating on-target mechanism. Additional evidence of cPheRS binding and inhibition in *T. gondii* was obtained via thermal-shift and aminoacylation assays, respectively (ΔTm 10.4 ºC, IC_50_ 0.42 μM). Of note, BRD2108 showed significantly weaker binding and inhibition of the human ortholog (ΔTm 3.5 ºC, IC_50_ 16 μM), suggesting high selectivity for the parasite target. Despite loss of activity with respect to the bicyclic azetidine lead BRD7929 (*Tg* growth EC_50_ 0.053 μM, *Tg*cPheRS ΔTm 11.6 ºC, aminoacylation IC_50_ 0.035 μM, Table 1),^9^ we deemed the bicyclic pyrrolidine series worthy of further investigation.

**Table 1.** Inhibition of *Tg*cPheRS mediates the antiparasitic activity of bicyclic pyrrolidines.

|  | BRD7929 | BRD2108 | mCMY416 |
| --- | --- | --- | --- |
| *Tg*RH Fluc EC<sub>50</sub><sup>a</sup> | 0.05<sup>c</sup> | 0.27 | 0.20 |
| *Tg*L497I Fluc EC<sub>50</sub><sup>a</sup> | 1.17<sup>c</sup> | 3.98 | 4.98 |
| *Tg* cPheRS ΔTm<sup>b</sup> | 11.6 | 10.4 | 10.5 |
| Hs cPheRS ΔTm<sup>b</sup> | 4.5 | 3.5 | 3.6 |
| *Tg* cPhe RS IC<sub>50</sub><sup>a</sup> | 0.04<sup>c</sup> | 0.42 | 0.24 |
| Hs cPheRS IC<sub>50</sub><sup>a</sup> | 18.00 | 15.58 | 3.58 |

<sup>a</sup> values in μM, average of 2-3 biological with 2-3 technical replicates.
<sup>b</sup> difference in melting temperature (Tm, $^{\circ}$ C), average of 3 biological replicates.
<sup>c</sup> values originally reported in Radke et al. (ref. 9)

Next, we turned to the optimization of BRD2108. In addition to improving antiparasitic activity, we recognized the importance for prospective toxoplasmosis therapeutics to achieve high levels of brain exposure. To help identify small molecules potentially capable of penetrating the blood-brain barrier (BBB), we systematically calculated central nervous system multiparameter optimization (CNS MPO)^15, 16^ scores. The CNS MPO score indeed weighs key physicochemical properties to predict a small molecule’s ability to penetrate the BBB. Scores can vary between 0 and 6.0, with higher scores indicating a higher likelihood of BBB penetration.

Early SAR studies focused on introducing major changes to the BRD2108 scaffold (Figure 2). kCMY297 mCMY298, and mCMY634 were synthesized to test the importance of the linkage between the proximal and distal ring systems. Unfortunately, these compounds were inactive. It appears that the alkyne imparts essential rigidity to the system and extends the distal ring into an active position. Lactams mCMY367 and mCMY371 lack the urea appendage that is expected, based on *Plasmodium* structural data,^12, 13^ to interact with the target by occupying the so-called auxiliary site; consistent with this model, these compounds were found to be inactive. *Trans* ring systems were also synthesized but the potency severely decreased in each case. BRD7929 is a 4,8-bicyclic azetidine that was an early lead in *T. gondii* studies from the HTS at the Broad Institute.^9^ This compound has good potency and exposure, but toxicity-associated liabilities. The PK of this and other compounds will be discussed later in this paper. It is clear that the alkyne linker, urea segment, and stereochemical configuration of the core are essential for potency against *T. gondii*. These requirements are consistent with crystallographic data of bicyclic azetidines bound to the active site of parasite PheRS enzymes: the biaryl alkyne group occupies the phenylalanine binding pocket while the methoxyphenyl that emanates from the urea occupies and auxiliary site that is unique to the parasite enzyme.

**Figure 2.**
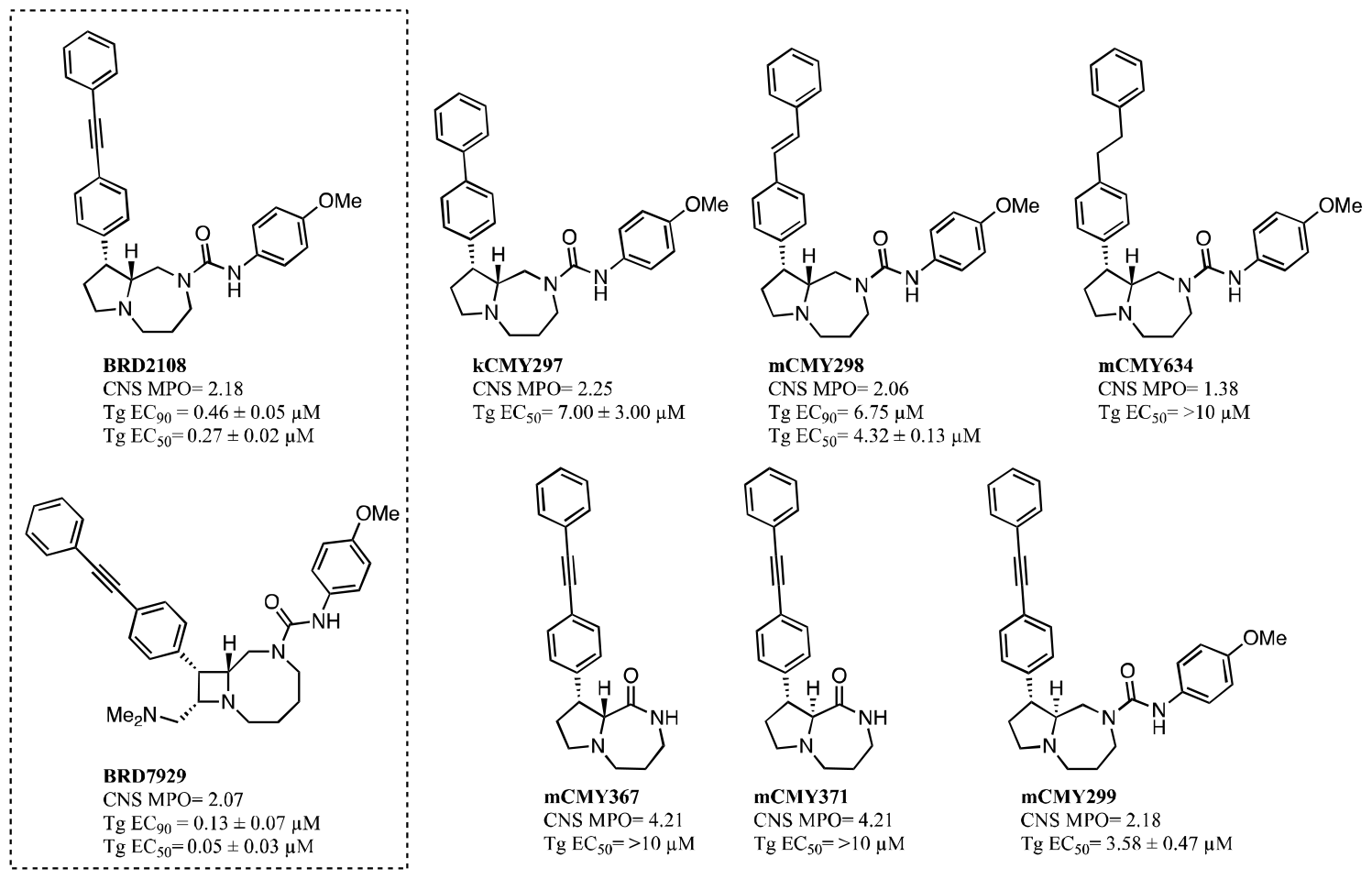
Early studies inform on key structural features of bicyclic pyrrolidine *Toxoplasma* growth inhibitors. Chemical structures of bicyclic pyrrolidine series analogs and of bicyclic azetidine lead BRD7929, and *in vitro T. gondii* inhibitory potencies. Potency values provided (Tg EC_50_, Tg EC_90_) are an average of n = 2-3 experiments (except mCMY414 where n = 1 and BRD7929 where n > 3); ± indicates standard deviation. EC_90_ was not determined for compounds with Tg EC_50_ > 3 μM.

Distal ring substituents of our targets could be explored with the core structure established, as shown in Table 2. We explored possible aniline (R_2_) substitutions on BRD 2108. Unfortunately, kCMY240, mCMY295, and mCMY369 did not reach the potency of BRD 2108. Replacement of R_2_ aromatic substituents with alicyclic substituents, as demonstrated by mCMY413, mCMY370, and mCMY373, also resulted in a stark decrease in potency.

A more extensive SAR study was conducted on the distal phenyl ring (R_1_, Table 2). Substituting R_1_ with heteroaromatic rings improved the CNS MPO score modestly but was detrimental to potency. For example, mCMY096, mCMY095, and mCMY174 are slightly active, with mCMY173 being inactive. mCMY049 and kCMY093 were inactive, while pyridazine (kCMY175) was only slightly active. mCMY176 and mCMY233 were synthesized to see if a smaller aliphatic group could be tolerated but both were found to be inactive. Potency suffered in both the *meta-*tolyl substitution in mCMY177 and the unconjugated system of mCMY232. Of note, the larger aliphatic system of mCMY230 maintained a similar potency to BRD2108. Previous reports based on testing of a limited set of bicyclic azetidines suggested that an aromatic distal ring in the biaryl alkyne was more potent than analogs bearing an aliphatic distal ring.^13^ However, sp^3^ systems introduce molecular complexity without significantly increasing molecular weight and can adjust molecular shape to optimize receptor/ligand compatibility. Aliphatic systems usually have higher selectivity and solubility compared to flat aromatic compounds and have been shown to correlate with more success in clinical trials.^17^ Therefore, we explored additional aliphatic rings not examined previously. While saturated systems mCMY412 and mCMY415 had modest potency, we were pleased to find that mCMY416 surpassed the potency of BRD2108. The higher potency of mCMY416 compared to mCMY412 (fully saturated) and BRD2108 (fully unsaturated) could be due to the conjugated system or the conformations sampled by the partially unsaturated ring. The smaller mCMZ070 was later synthesized but was not as active as mCMY416. This encouraging result prompted us to test other heteroatoms in these rings. The distal tetrahydropyran (mCMY544), difluoro (mCMY835), tetrahydrothiopyran (mCMZ157), and tetrahydropyran-ol (mCMZ235) had modest activity. On the other hand, propargyl (mCMY546) and alcohol mCMZ557 had potency comparable to mCMY416. mCMZ557 also has an increased CNS MPO score as well as two more chiral centers. This series of compounds shows that potency can be maintained with a diversity of saturated systems.

**Table 2.** Replacement of distal aromatic ring in BRD2108 with sp^3^-rich system preserves *in vitro* antiparasitic potency SAR of distal phenyl ring and aniline moiety with EC_90_ values. Distal heteroaromatic compounds lost potency but aliphatic compounds maintained potency and show diversity at distal position. *p-*tolyl and methoxy anilines had the best potency but aliphatic and smaller substituents lost potency. Antiparasitic potency values provided (Tg EC_50_, Tg EC_90_) are an average of n = 2-3 experiments unless otherwise noted; ± indicates standard deviation. Compounds for which standard deviation is not provided were tested in only 1 experiment or exceeded a 10-μM EC_90_.

| ID | R<sub>1</sub> | R<sub>2</sub> | EC<sub>90</sub> (μM) |
| --- | --- | --- | --- |
| BRD2108 |  |  | 0.46 ± 0.05 |
| mCMY096 |  |  | 4.88 |
| mCMY095 |  |  | 3.87 |
| mCMY173 |  |  | >10 |
| mCMY174 |  |  | 4.17 |
| mCNV579 |  |  | 0.96 ± 0.06 |
| mCMY049 |  |  | 9.69 |
| kCMY093 |  |  | >10 |
| kCMY175 |  |  | 3.63 ± 0.13 |
| mCMY176 |  |  | >10 |
| mCMY233 |  |  | >10 |
| mCMY230 |  |  | 0.71 ± 0.22 |
| mCMY177 |  |  | 5.60 |
| mCMY232 |  |  | >10 |
| kCMY240 |  |  | 4.82 ± 0.13 |
| mCMY295 |  |  | 1.26 ± 0.08 |
| mCMY369 |  |  | 5.42 |
| mCMY413 |  |  | 4.63 ± 0.08 |
| mCMY370 |  |  | >10 |
| mCMY373 |  |  | 4.78 |
| mCMY412 |  |  | 1.81 ± 0.41 |
| mCMY415 |  |  | 1.21 ± 0.09 |
| mCMY416 |  |  | 0.38 ± 0.02 |
| mCMZ070 |  |  | 3.09 ± 0.19 |
| mCMY544 |  |  | 3.07 ± 0.33 |
| mCMY546 |  |  | 0.46 ± 0.04 |
| mCMY835 |  |  | 3.89 ± 0.32 |
| mCMZ157 |  |  | 2.10 ± 1.10 |
| mCMZ557 |  |  | 0.52 ± 0.06 |
| mCMZ235 |  |  | 4.88 ± 0.73 |

Seven compounds exhibiting high antiparasitic potency (EC_50_ < 1 μM and EC_90_ < 1.5 μM) were selected for further profiling. The selectivity, cross-resistance, and ADME profiles of these compounds are shown in Table 3. Among these, BRD2108 had average selectivity in the THP-1 (7.94 μM) and HepG2 (8.48 μM) assays in relation to the other compounds tested in this series. BRD2108 had the best microsomal stability and MDCK-MDR1 permeability with a B-A/A-B ratio of 2.84. Unfortunately, like most other compounds of this series the plasma protein binding proved very high. mCMY230 was an unremarkable compound in terms of potency, selectivity, CNS MPO and AlogD score, plasma protein binding, and microsomal stability. mCMY295 was slightly less potent but has the best CNS MPO (2.45) and AlogD (3.4) score due to the distal pyridine ring. mCMZ557 had a lower PPB and BHB than any other compound. mCMY546 had slightly lower PPB and BHB than the average, but the highest microsomal stability of all compounds. mCMY416 is the most potent compound synthesized but has the lowest selectivity (3.14 μM in THP-1 and 2.24 μM in HepG2).

**Table 3.**
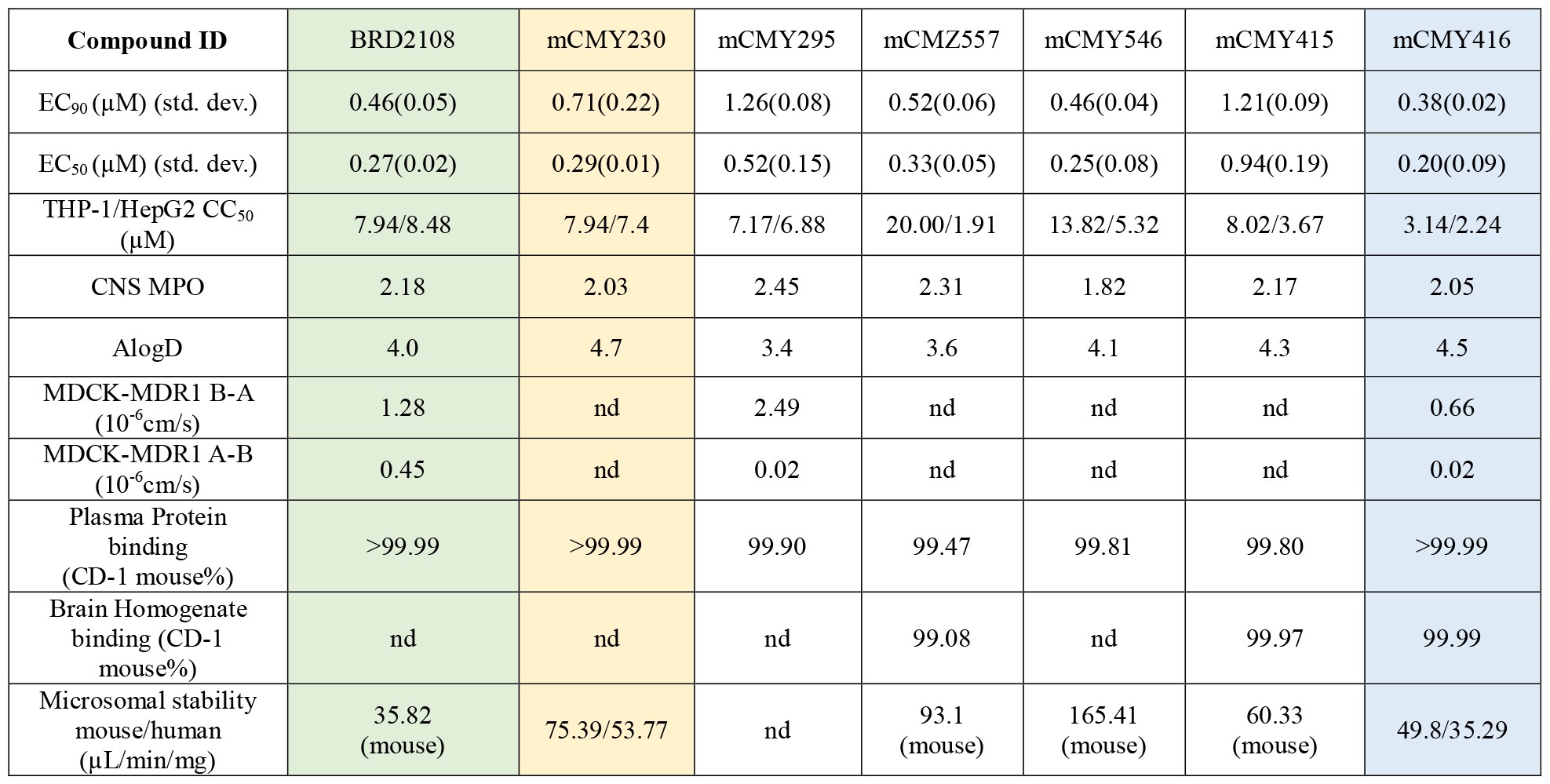
ADME, selectivity, and cross-resistance profiles for lead compounds BRD2108, mCMY416, and potent bicyclic pyrrolidine congeners. nd: not determined

The pharmacokinetic (PK) data for BRD2108, mCMY416, mCMY546 and BRD7929 are shown in Table 4. These PK experiments were run in solution at 10mg/kg in mice. BRD2108 has the best PK data with very good brain exposure that reaches 10X the EC_90_ at 8 hours with 3042 ng/g. Plasma exposure is not as high (325ng/mL) but does cover the EC_90_. BRD2108 also has the best B/P ratio of all three compounds. mCMY416 does not have a high brain exposure, but still covers the EC_90_ by 2X (C_max_ is 339ng/g at 1 hour). mCMY546 has much lower exposure and a higher EC_90_. As a result, the EC_90_ is not covered by the brain exposure. mCMY546 also has the lowest B/P ratio of the three compounds. The PK of BRD7929 shows that the EC_90_ is well covered by brain exposure (1790ng/g at 8 hours). While mCMY416 and mCMY546 start to clear after 1 hour and BRD2108 levels out after 3 hours, the BRD7929 exposure increases steadily until 8 hours. This poses a risk of accumulation and may be a challenge to dose safely. mCMY416 has a more typical drug PK profile than BRD7929 where EC_90_ is covered in the brain and begins to clear after a few hours.

**Table 4.**
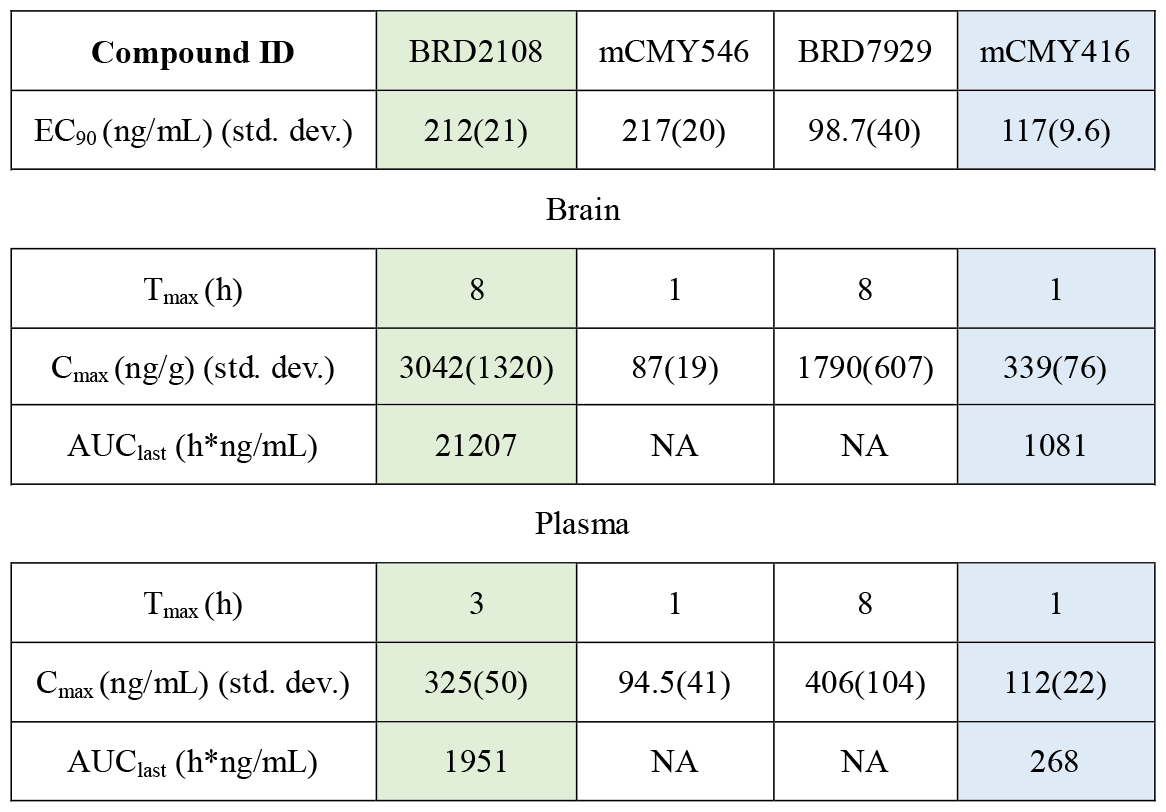
Pharmacokinetic data for compounds BRD2108, mCMY416, mCMY546, and BRD7929. CD-1 mice (n = 6 per treatment arm), 10 mg/kg p.o. single dose, formulation: 75%PEG300:25%D5W (solution).

| Compound ID | BRD2108 | mCMY546 | BRD7929 | mCMY416 |
| --- | --- | --- | --- | --- |
| EC<sub>90</sub> (ng/mL) (std. dev.) | 212(21) | 217(20) | 98.7(40) | 117(9.6) |
| Brain |  |  |  |  |
| T<sub>max</sub> (h) | 8 | 1 | 8 | 1 |
| C<sub>max</sub> (ng/g) (std. dev.) | 3042(1320) | 87(19) | 1790(607) | 339(76) |
| AUC<sub>last</sub> (h*ng/mL) | 21207 | NA | NA | 1081 |
| Plasma |  |  |  |  |
| T<sub>max</sub> (h) | 3 | 1 | 8 | 1 |
| C<sub>max</sub> (ng/mL) (std. dev.) | 325(50) | 94.5(41) | 406(104) | 112(22) |
| AUC<sub>last</sub> (h*ng/mL) | 1951 | NA | NA | 268 |

To better understand the PK profile of mCMY416, a higher dose (100 mg/kg) in suspension was administered. Table 5 shows a comparison between the 10 mg/kg solution dosing vs. the 100 mg/kg suspension dosing. In this experiment, brain exposure was much higher due to a delayed absorption of the compound. At 8 hours, the concentration in the brain reached 15,400 ng/g which covers the EC_90_ (177 ± 9.6 ng/mL) by more than 80X. In a later PK study, time points were taken from 8-48 hours to determine the clearance and the risk for accumulation. The compound started to clear at around 20 hours. The suspension dosing must have better absorption compared to the solution dosing because the concentration for 100 mg/kg was over-proportional in the brain at 45X higher than the 10mg/kg dosing and the AUC is 55X larger in the 100mg/kg dosing than the 10 mg/kg dosing. The plasma C_max_ was closer to expected with 100 mg/kg dosing reaching 1,584 ng/mL and 10 mg/kg reaching 112 ng/mL (about 14X).

**Table 5.**
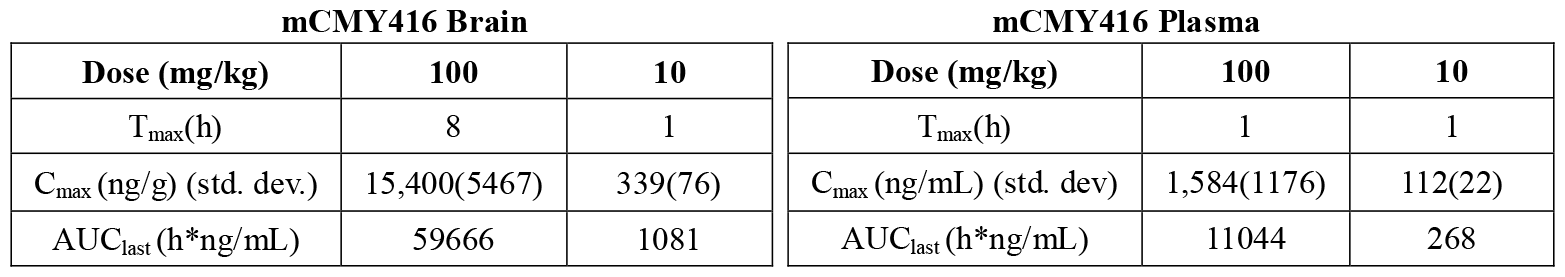
Pharmacokinetic data for compound mCMY416: comparison between 100 mg/kg suspension dose and 10 mg/kg solution dose. CD-1 mice (n = 6 per treatment arm), 10 mg/kg single dose p.o. (formulation: 75%PEG300:25%D5W [solution]), 100 mg/kg single dose p.o. (0.2% MC/Tween [suspension]).

| mCMY416 Brain |  |  | mCMY416 Plasma |  |  |
| --- | --- | --- | --- | --- | --- |
| Dose (mg/kg) | 100 | 10 | Dose (mg/kg) | 100 | 10 |
| T<sub>max</sub> (h) | 8 | 1 | T<sub>max</sub> (h) | 1 | 1 |
| C<sub>max</sub> (ng/g) (std. dev.) | 15,400(5467) | 339(76) | C<sub>max</sub> (ng/mL) (std. dev.) | 1,584(1176) | 112(22) |
| AUC<sub>last</sub> (h*ng/mL) | 59666 | 1081 | AUC<sub>last</sub> (h*ng/mL) | 11044 | 268 |

In conclusion, we have identified a series of bicyclic pyrrolidine inhibitors of the parasite causative of toxoplasmosis act by inhibiting the parasite phenylalanyl t-RNA synthetase (TgcPheRS). The bicyclic pyrrolidine core scaffold, typified by the small molecule BRD2108, was designed to retain the exit vector geometry of the bicyclic azetidine isomeric series, typified by BRD7929 and known for activity against the same target enzyme. SAR investigation into BRD2108, which included variations to both core scaffold and appendages, led us to the discovery of a more potent compound, mCMY416. mCMY416 integrates sp^3^ carbons into the distal ring and creates opportunities for diversification at this position. Pharmacokinetic studies in mouse show that oral dosing of mCMY416 leads to compound concentrations in the brain that exceed *in vitro* EC_90_, holding promise for achieving antiparasitic efficacy *in vivo*. Taken together, pharmacokinetic and biological assays conducted on the most potent compounds identified so far also suggest potential for improvement in permeability and free fraction in the brain. The discovery of mCMY416, which provides opportunities for accessing further chemical diversity in the PheRS inhibitor space, opens an avenue for improvement in these areas.

## Experimental Section

### 1. Synthesis

#### General Considerations

Reagents and solvents were commercially obtained and used without further purification. Anhydrous solvents were used and reactions performed under atmosphere of Argon. Crude products were purified on silica gel chromatography or C18 reverse-phase columns. ^1^H NMR spectra were obtained on Bruker Ultrashield 400 MHz. Shifts are expressed in ppm (parts per million) and coupling constants (*J*) in Hertz (Hz). LC-MS was obtained on a Waters Acquity Ultra Performance LC system.

#### Synthetic Route

The synthetic route to mCMY416 and other leads is described in Scheme 1. Synthesis begins with the HATU coupling of commercially available Fmoc-D-Pro-OH (**1**) and the directing group 8-Aminquinoine in almost quantitative yield. Intermediate **2** is then used in the Palladium(II) acetate catalyzed C-H activation to form intermediate **3**. Although conversion and yield in this reaction is modest, the aminoquinoline directing group gives almost exclusively the *cis* conformation. After quantitative deprotection to form **4**, the directing group removal can be performed using 18%HCl in H_2_O solution to give intermediate **5**. Racemization is observed from 10-50% depending on conditions and the scale of the reaction. *Trans* isomers could be used to synthesize the *trans* compounds in Figure 2. Reductive amination using NaBH(OAc)_3_ and the Fmoc-protected aldehyde, **6** adds the carbon chain that will become the 7-membered ring. One-pot Fmoc removal using Silia-Bond® piperazine and subsequent intramolecular ring formation gives lactam **8** at a good yield of 75%. Ruthenium-catalyzed reduction of the lactam using TMDS as the hydride source and then subsequent isocyanate addition gives urea **9** (80% yield when R = OMe). **9** can then be used in all SAR described in this paper through a Sonogashira-like coupling with the desired alkyne with the aryl bromide in good yield.

**Scheme 1.**
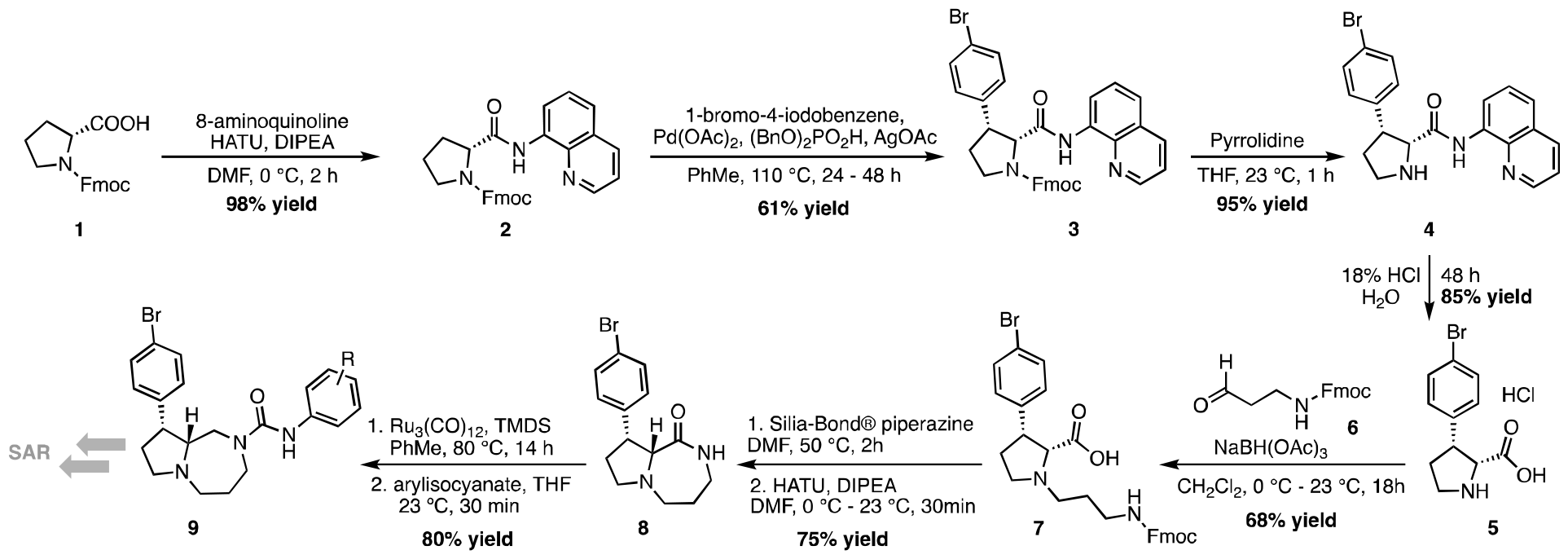
Synthetic route used for the preparation of mCMY416 and analogs.

#### (9*H*-fluoren-9-yl)methyl (*R*)-2-(quinolin-8-ylcarbamoyl)pyrrolidine-1-carboxylate (2)

Fmoc-D-Proline-OH (**1**) (10 g, 29.7 mmol) was dissolved in DMF (75 mL) and cooled to 0 °C. HATU (11.3 g, 1 eq), DIPEA (10.3 mL, 2 eq), and 8-aminoquinoline (21.3 g, 5 eq) were then added sequentially. The reaction was stirred at 0 °C for 2 hours. Sat. aq. NH_4_Cl was added and the mixture was extracted with EtOAc. The organic layer was washed with 3M HCl until all excess 8-aminoquinoline was removed. (The next reaction will not proceed efficiently if 8-aminoquinoline is present). The organic layer was then washed with saturated aqueous NaHCO_3_ until color changes to white. The organic layer was then washed with brine and then dried with Na_2_SO_4_, filtered, and concentrated to afford a white solid product, which was used in the next step without further purification (98% yield, 13.4 g, 29 mmol).

#### (9*H*-fluoren-9-yl)methyl (2*R*,3*R*)-3-(4-bromophenyl)-2-(quinolin-8-ylcarbamoyl)pyrrolidine-1-carboxylate (3)

Compound **2** (13.8 g, 29.7 mmol), Pd(OAc)_2_ (667 mg, 0.1 eq), dibenzyl phosphate (1.65 g, 0.2 eq), AgOAc (9.91 g, 2 eq), and 1-Bromo-4-Iodobenzene (25.1 g, 3 eq) were purged with argon and then dissolved in dry toluene (300mL) with 3-Å molecular sieves. The reaction was heated to reflux for 24-48 hours under argon and then concentrated onto Celite for purification by column chromatography (1:1 Hexane/Ethyl Acetate), affording 8.3 g of the title product as a yellow solid (61% yield, 11.2 g, 18.1 mmol).

#### (2*R*,3*R*)-3-(4-bromophenyl)-*N*-(quinolin-8-yl)pyrrolidine-2-carboxamide (4)

Compound **3** (8.3 g, 13.4 mmol) was dissolved in dry THF (67 mL). Pyrrolidine (11 mL, 10 eq) was then added dropwise and the reaction was stirred for 2 hours at room temperature. The reaction was then concentrated and purified by column chromatography (1:1 Hexane/EtOAc to EtOAc, then DCM to 8:2 DCM/MeOH) to afford the title product as a yellow solid (95% yield, 5.03 g, 12.7 mmol).

#### (2*R*,3*R*)-3-(4-bromophenyl)pyrrolidine-2-carboxylic acid (5)

Compound **4** (5.3 g, 13.4 mmol) was dissolved in 290 mL of 17% HCl_(aq)_ and heated in a sealed vial at 115 °C for 17 hours (note: too concentrated acid or heating too long/hot will cause isomerization to *trans* compound; in ^1^H NMR *cis* compound has doublet at 4.65 ppm in MeOD but *trans* has doublet at 4.41 ppm in MeOD). The reaction was cooled and the water was removed under reduced pressure. Purification option 1) Water was added and the compound was dried on the lyophilizer. Solids were then dissolved in 7:3 CHCl_3_/iPrOH and filtered. The liquid was concentrated to give a yellow oil and used in the next step. Purification option 2) After the water was removed after reaction, aqueous NaOH was added and washed with EtOAc to remove any starting material. The mixture was acidified with aqueous HCl to pH 2 and water was removed. 7:3 CHCl_3_:iPrOH can be used dissolve product to remove salts. (85% yield, 3.46 g,11.4 mmol of yellow oil-purification option 1). Diastereomers can be separated after lactam formation.

#### (9*H*-fluoren-9-yl)methyl (3-oxopropyl)carbamate (6)

To a solution of Fmoc-β-alaninol (5.8 g, 2 eq) in dry DCM (60 mL) at 0 °C was added Dess-Martin Periodinane (12.5 g, 3 eq). The reaction was allowed to warm to room temperature and stirred for about 4 hours. Saturated aqueous NaHCO_3_ (200 mL) and Et_2_O (200 mL) were added and the mixture was stirred vigorously for 10 minutes. After filtering off solids, the layers were separated and the aqueous layer was extracted 3X with Et_2_O. The organic layer was washed with brine and dried over Na_2_SO_4_. After evaporation of the solvent (below 30 °C to prevent polymerization) the aldehyde (white solid) was used in the next step without further purification.

#### (2*R*,3*R*)-1-(3-((((9*H*-fluoren-9-yl)methoxy)carbonyl)amino)propyl)-3-(4-bromophenyl)pyrrolidine-2-carboxylic acid (7)

Compound **5** (3 g,9.87 mmol) was dissolved in dry DCM (50 mL) and NaBH(OAc)_3_ (6.25 g, 3 eq) was added at 0 °C while stirring. Aldehyde **6** was dissolved in dry DCM (40 mL) and added dropwise to compound **5** at 0 °C. The reaction was warmed to room temperature and stirred overnight. The reaction was then quenched with NH_4_Cl and extracted with DCM. The product was purified by column chromatography (1:1 Hexane/EtOAc to EtOAc, then DCM to 8:2 DCM/MeOH) to give the title product as a white solid (68% yield, 3.68 g, 6.7 mmol).

#### (9*R*,9a*R*)-9-(4-bromophenyl)octahydro-1*H*-pyrrolo[1,2-a][1,4]diazepine (8)

Compound **7** (2.6 g, 4.74 mmol) was dissolved in dry DMF (475 mL). *SiliaBond*^®^ *Piperazine* (16.9 g, 3 eq) was added and heated to 50 °C for 2 hours. The reaction was then cooled to 0 °C and HATU (2.7 g, 1.5 eq) and DIPEA (4.1 mL, 5 eq) were added. The reaction was then warmed to room temperature and stirred for 20 minutes. DCM and SiO_2_ were added and concentrated. The solids were transferred to fritted funnel and washed with hexanes and the liquids were discarded. The solids were then rinsed with 20% MeOH in DCM and the liquids were concentrated. EtOAc was then added and washed with saturated aqueous NaHCO_3_ and dried over Na_2_SO_4_ and then concentrated. The residue was purified by column chromatography (1:1 Hexane/EtOAc to EtOAc, then DCM to 8:2 DCM/MeOH) to give the title product as a yellow oil (75% yield, 1.09 g, 3.56 mmol). (*Trans* isomer elutes slightly after *cis* isomer.)

#### (9*R*,9a*R*)-9-(4-bromophenyl)-*N*-(4-methoxyphenyl)hexahydro-1*H*-pyrrolo[1,2-a][1,4]diazepine-2(3*H*)-carboxamide (9 [R = OMe])

Compound **8** (666 mg, 2.16 mmol) was added to a flame-dried vial along with Ru_3_(CO)_12_ (207 mg, 0.15 eq) and flushed with argon. Dry toluene (1.35 mL) was added and vortexed for 30 seconds. TMDS (5.7 mL, 15 eq) was added and the vial was sealed and heated to 80 °C for 6 hours. Reaction was cooled to room temperature and concentrated. Crude residue was dissolved in dry THF (8.6 mL) and 4-methoxyphenyl isocyanate (0.28 mL, 1 eq) was added slowly at 0 °C and then stirred one hour at room temperature. The reaction was quenched with MeOH and then concentrated. The crude mixture was purified by column chromatography (9:1 DCM/MeOH) to afford the title product as a brown solid (75% yield, 720 mg recovered, 1.57 mmol).

#### 9*R*,9a*R*)-9-(4-(cyclohex-1-en-1-ylethynyl)phenyl)-*N*-(4-methoxyphenyl)hexahydro-1*H*-pyrrolo[1,2-a][1,4]diazepine-2(3*H*)-carboxamide) (mCMY416)

Compound **9 [R = OMe]** (150mg, 0.34mmol), 1-ethynylcyclohex-1-ene (0.158 mL, 4 eq), and XPhos-Pd-G3 (29 mg, 0.1 eq), were added to a dry vial and flushed with argon. Acetonitrile (1.7mL) was sparged with argon and added to solid reagents and then triethylamine (0.19 mL, 4 eq) was added. The reaction was stirred at 70 °C overnight. The reaction was cooled to room temperature and then quenched with saturated aqueous NaHCO_3_. The layers were separated, the organics were extracted with EtOAc, then dried over Na_2_SO_4_. The product was purified by column chromatography (DCM to 95:5 DCM/MeOH) followed by ISCO reverse column to afford the title product as a white solid (57% yield, 91 mg, 0.19 mmol).

### 2. DMPK Assays

#### Microsomal stability and plasma protein binding studies

Studies were carried out at WuXi AppTec (Shanghai, China). For LC/MS/MS analysis, test and control compounds were quantified using peak area ratio of analyte and internal standard. The metabolic stability in liver microsomes was determined using the compound depletion approach and quantified by LC/MS/MS. Mouse microsomes were used (BD Gentest). The rate and extent of metabolism was determined by the disappearance of the parent compound, allowing for the determination of in vitro half-life (t1/2), intrinsic clearance (Clint) and the extraction ratio (ER) in various species. Protein binding in plasma was determined using equilibrium dialysis. Compounds were tested at 2 μM in triplicate, and concentrations were quantified by LC/MS/MS.

### 3. *In vivo* assays

#### Animal Study: Animal Care and Compound Application

Animal experimental procedures were approved by the Institutional Animal Care and Use Committee (IACUC) of the respective study locations. Pharmacokinetic studies were conducted at WuXi (China).

#### Pharmacokinetics in mouse

For all (IV or PO) reported pharmacokinetic studies in this manuscript, three fasted animals per study group were administered the test article as a solution in Formulation:75% PEG300:25%D5W for solution or Formulation: 0.2% MC/Tween for suspension. Blood and brain samples were collected at 0.5, 1, 3, 5, 8, 24 hr post-dosing for PO studies.

The blood samples were centrifuged to obtain the plasma, plasma was stored at below -20°C until analysis. Plasma concentrations were determined by liquid chromatography/tandem mass spectrometry (LC−MS/MS). The PK parameters were determined by non-compartmental methods using WinNonLin (v6.1 or higher version, Certara Inc.).

### 4. *In vitro* Assays for Growth and Cytotoxicity

Two strains of *Toxoplasma gondii*, namely TgRH-FLuc (type I)^18^ and TgRH-PheRS^[L497I]-^FLuc^9^, were previously documented. Tachyzoites stage of these parasite lines were cultured through sequential passages in confluent human foreskin fibroblasts (HFF). Both HFF and parasite cultures were nurtured in a medium consisting of 10% DMEM (Dulbecco’s Modified Eagle Medium), supplemented with 10 mM glutamine, 10 μg/mL gentamycin, and 10% fetal bovine serum. The cultures were incubated at 37°C with 5% CO_2_ and routinely checked for mycoplasma contamination using the e-Myco Plus kit from Intron Biotechnology. Human hepatocellular carcinoma cells (HepG2, ATCC-HB-8065) and the human monocytic tumor line (THP-1, ATCC-TIB-202) were employed to assess toxicity against host cells. These cell lines were maintained according to the specifications recommended by ATCC.

The compounds utilized were obtained from Calibr, provided either as lyophilized powder or as 10 mM stocks in 100% DMSO, stored at −80°C before use. Luciferase-based growth assays were conducted in the inner 60 wells of a clear-bottom, white 96-well plate (Costar) to prevent inter-well interference^16^. The compounds were diluted to create a 3-fold dilution series. Each well received freshly harvested parasites (5×10^3^) in a 100 μL volume to achieve a final compound concentration (200 μL/well total volume, 0.1% DMSO in 10% DMEM medium). Incubation of plates occurred for 72 hours at 37°C before analysis, adhering to the Luciferase Assay System protocol from Promega^9, 18, 19^. Post-incubation, the culture media were replaced with 50 μL of 1x Cell Culture Lysis Reagent (Promega) for 10 minutes at room temperature. Subsequently, 100 μL of Luciferase Assay reagent was added to each well, and luciferase activity was measured using a BioTek Cytation 3 multi-mode imager equipped with Gen5 software (v3.08).

For toxicity screening, compounds were diluted to 120 μM (2x concentration in 10% DMEM, 0.4% DMSO) before creating a 10-dose series through step-wise, 3-fold dilutions in 10% DMEM. Host cells were seeded at a density of 10x10^3^ cells/well (100 μL vol), facilitating sub-confluent monolayers to allow expansion during the 3-day growth assay. THP-1 cells underwent treatment with 10 ng/ml phorbol 12-myristate 13 acetate (PMA) for 24 hours to differentiate into macrophages before compound addition. In the case of HepG2 cells, compounds were added 6 hours post-seeding into host cell-containing plates (200 μL final volume, 0.2% DMSO) and incubated in culture medium at 37°C supplemented with 5% CO2. At 72 hours post compound addition, the culture media were aspirated to 80ul, added same volume of CellTiter-Glo® Luminescent Cell Viability Assay reagent (Promega), and luciferase activity was measured using a BioTek Cytation 3 quipped with Gen5 software (v3.08).

All assays were repeated with two technical replicates within each of two or three independent biological replicates. Statistical analyses were performed using Prism 10 (GraphPad Software, Inc.). Dose–response inhibition curves for parasite and host cell toxicity screens (EC50 values) were generated using the “Log(inhibitor) vs. normalized response—Variable slope” function. Values reported are means ± standard deviation.

### 5. Protein Expression and Purification

The full-length cytosolic parasitic (*Tg*cPheRS) and host (*Hs*cPheRS) genes were synthesized and codon optimized for efficient expression in *E. coli* as described earlier^9, 12^. The genes encoding alpha and beta subunit of *Tg*cPheRS enzyme were fused. A similar approach was employed for *Hs*cPheRS enzyme. These composite genes were then cloned in pETM11 using Nco1 and Kpn1 restriction enzymes. The transformation process was carried out in BL-21 strain of *E. coli* for both the constructs. Briefly, the *E. coli* cultures were cultivated at 37 □°C until reaching an OD_600_ of 0.6–0.8. Induction of recombinant protein expression was done by the addition of 0.6 □mM isopropyl β-d-1-thiogalactopyranoside (IPTG) at 18 □°C. Following, 18–20 □h post-induction period, the cells were harvested via centrifugation at 5,000 □g for 20 □min, resuspended in binding buffer (50 □mM Tris–HCl (pH 8), 200 □mM NaCl, 4 □mM β-mercaptoethanol (βMe), 10% (v/v) glycerol, 1 □mM phenylmethylsulfonyl fluoride (PMSF) and 0.1 □mg/mL lysozyme), and lysed using sonication. The lysate underwent a subsequent centrifugation at 20,000 □g for 45 □min and the cleared soluble protein lysate was loaded on a prepacked Ni-NTA column (GE Healthcare). Proteins binding to the Ni-NTA coloumns were washed with 20 column volumes of binding buffer supplemented with 20 □mM imidazole to eliminate impurities. Elution of bound proteins occurred using a concentration gradient of imidazole (0 to 1 □M) in elution buffer (80 □mM NaCl, 50 □mM Tris–HCl (pH 8), 4 □mM β-mercaptoethanol, 10% (v/v) glycerol, 1 □M imidazole, using AKTA-FPLC system (GE healthcare)). The fractions containing the purified protein were pooled together and concentrated using 30-kDa cut-off centrifugal devices (Millipore). Further, size exclusion chromatography using the GE HiLoad 60/600 Superdex column was performed in a buffer comprising 50 □mM HEPES (pH 8), 200 □mM NaCl, 4 □mM β-mercaptoethanol, 1 □mM MgCl_2._ Purity of the eluted fractions for both the proteins was confirmed via SDS PAGE gel, followed by pooling and storage of fractions −80 □°C until further use.

#### Thermal Shift Assays

These were performed with *Tg*cPheRS and *Hs*cPheRS enzymes with BRD7929, BRD2108, mCMY416 by diluting the purified enzyme to a final concentration of 2 μM in a buffer containing 200 mM NaCl, 5 mM MgCl2, 50 mM Tris (pH 7.5), and 2 X SYPRO orange dye (Life Technologies)^6, 12, 13^. Each compound at a concentration of 20 μM was individually added to the enzyme solutions and samples were incubated at room temperature for 10min. The solution containing the purified enzymes alone (Apo) and with natural substrates (ATP and L-phenylalanine) were kept as control. The samples were briefly subjected to temperature ramp from 25°C TO 99°C at a rate of 1°C min^-1^ and the fluorescence signals of SYPRO® orange dye were monitored using StepOnePlus ™ quantitative real-time PCR system (Life Technologies). Each melting curve is an average of three measurements and analysis was done using Protein Thermal shift software (v1.3, Thermofisher). The derivative Tm was used for analysis.

#### Enzyme Assays

The enzyme inhibition assays with *Tg*cPheRS and *Hs*cPheRS were performed using malachite green dye based aminoacylation assays as previously described.^15,17^ The assays were carried out at 100nM of purified enzyme concentrations in presence of 50 μM L-phenylalanine and 100 □μM ATP. The reaction was done in a standard aminoacylation buffer comprising 30 □mM HEPES (pH 7.5), 50 □mM MgCl_2,_ 150 □mM NaCl, 30 □mM KCl, 1 □mM DTT. The assays were initiated by adding 2 □U/mL *E. coli* inorganic pyrophosphatase enzyme. For the determination of IC_50_ values, a 10-fold dilution of the inhibitors was done in 96-well clear, flat bottom plates with starting concentration of 25uM. The complete reaction mixtures were incubated at 37°C for 2hours. The reactions were terminated using malachite green stop solution and activity of enzymes was measured at 620 □nm using SpectraMax M2 (Molecular Devices). MBP (maltose binding protein) was used a negative control for protein binding the reaction mixture. All the experiments were repeated three independent times with internal triplicates. The statistical analysis was performed using graph pad Prism6.

## Acknowledgements

This work was supported by a grant from the National Institutes of Health (AI143857). A.S. is supported by a JC Bose fellowship.

## Declarations

The authors declare there are no conflicts to report.

## List of abbreviations

Tg: Toxoplasma gondii
SAR: structure-activity relationship
PheRS: phenylalanine tRNA synthetase
BBB: blood brain barrier
CNS MPO: central nervous system multiparameter optimization
PK: pharmacokinetic
PEG: polyethylene glycol
D5W: Dextrose 5% in water
T_max_: time at maximum concentration
C_max_: Maximum concentration
AUC_last_: Area under the curve from time of dosing to time of last observation
PPB: plasma protein binding
BHB: brain homogenate binding

