## Supporting information for "Bicyclic pyrrolidine inhibitors of *Toxoplasma gondii* phenylalanine t-RNA synthetase with antiparasitic potency *in vitro* and brain exposure"

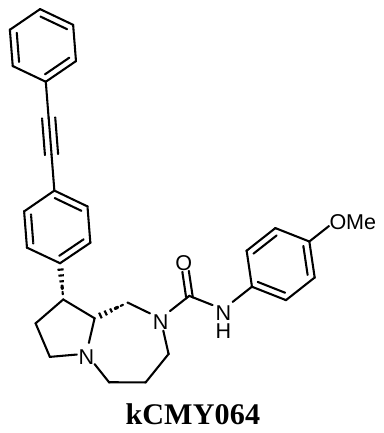

(9R,9aR)-N-(4-methoxyphenyl)-9-(4-(phenylethynyl)phenyl)hexahydro-1H-pyrrolo[1,2-a][1,4]diazepine-2(3H)-carboxamide: ^1^H NMR (400 MHz, MeOD) δ 7.50 – 7.45 (m, 2H), 7.44 – 7.40 (m, 2H), 7.38 – 7.32 (m, 5H), 7.16 (d, *J* = 9.3 Hz, 2H), 6.80 (d, *J* = 9.3 Hz, 2H), 3.73 – 3.72 (m, 3H), 3.59 (s, 2H), 3.52 – 3.45 (m, 1H), 3.42 (s, 1H), 3.23 – 3.14 (m, 2H), 2.65 (t, *J* = 9.3 Hz, 1H), 2.45 (d, *J* = 17.2 Hz, 2H), 2.30 – 2.21 (m, 2H), 2.03 – 1.92 (m, 3H). LC-MS *m/z* calcd. C_30_H_32_N_3_O_2_^+^ [M+H]^+^ = 466.62495, found = 466.4111

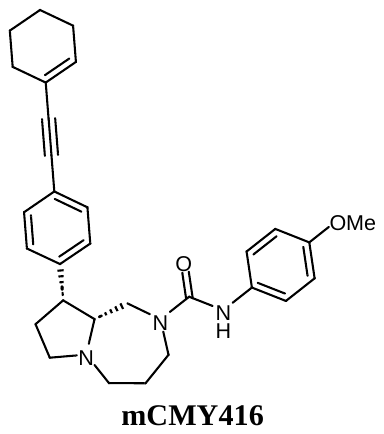

(9R,9aR)-9-(4-(cyclohex-1-en-1-ylethynyl)phenyl)-N-(4-methoxyphenyl)hexahydro-1H-pyrrolo[1,2-a][1,4]diazepine-2(3H)-carboxamide:^1^H NMR (400 MHz, Acetone) δ 7.43 (s, 1H), 7.40 – 7.30 (m, 6H), 6.79 (d, *J* = 9.2 Hz, 2H), 6.17 – 6.12 (m, 1H), 3.81 – 3.69 (m, 5H), 3.43 – 3.34 (m, 2H), 3.25 – 3.14 (m, 2H), 2.56 (t, *J* = 8.8 Hz, 1H), 2.41 – 2.28 (m, 3H), 2.23 – 2.13 (m, 5H), 1.97 – 1.90 (m, 3H), 1.70 – 1.60 (m, 4H). LC-MS *m/z* calcd. C_30_H_36_N_3_O_2_^+^ [M+H]^+^ = 470.6370, found = 470.6835.

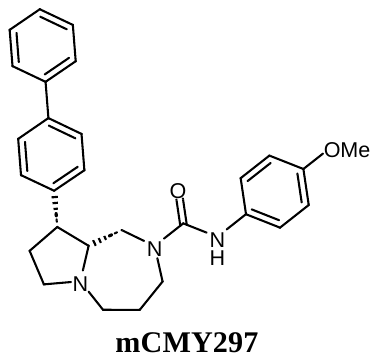

(9R,9aR)-9-([1,1'-biphenyl]-4-yl)-N-(4-methoxyphenyl)hexahydro-1H-pyrrolo[1,2-a][1,4]diazepine-2(3H)-carboxamide: ^1^H NMR (400 MHz, Acetone) δ 7.67 (d, *J* = 7.9 Hz, 2H), 7.60 (d, *J* = 8.3 Hz, 2H), 7.50 – 7.42 (m, 5H), 7.39 (d, *J* = 9.3 Hz, 2H), 7.36 (d, *J* = 7.3 Hz, 1H), 6.81 – 6.77 (m, 2H), 3.83 (d, *J* = 14.9 Hz, 1H), 3.74 (s, 4H), 3.48 – 3.35 (m, 2H), 3.25 (t, *J* = 7.9 Hz, 1H), 3.18 (dd, *J* = 12.8, 4.5 Hz, 1H), 2.62 – 2.56 (m, 1H), 2.44 – 2.37 (m, 2H), 2.35 – 2.20 (m, 2H), 2.01 – 1.90 (m, 3H). LC-MS *m/z* calcd. C_28_H_32_N_3_O_2_^+^ [M+H]^+^ = 442.2495, found = 422.3695

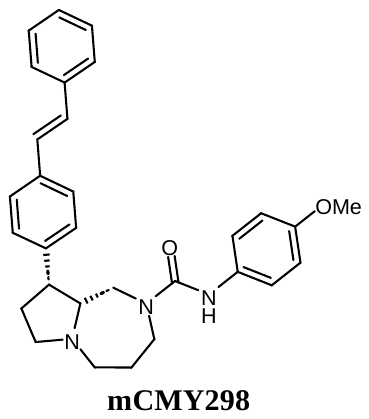

(9R,9aR)-N-(4-methoxyphenyl)-9-(4-((E)-styryl)phenyl)hexahydro-1H-pyrrolo[1,2-a][1,4]diazepine-2(3H)-carboxamide: ^1^H NMR (400 MHz, Acetone) δ 7.60 (d, *J* = 8.1 Hz, 2H), 7.53 (d, *J* = 8.3 Hz, 2H), 7.46 – 7.34 (m, 7H), 7.30 – 7.23 (m, 3H), 6.82 – 6.76 (m, 2H), 3.81 (d, *J* = 14.1 Hz, 1H), 3.74 (s, 4H), 3.46 – 3.34 (m, 2H), 3.24 (d, *J* = 8.4 Hz, 1H), 3.18 (dd, *J* = 12.6, 4.2 Hz, 1H), 2.58 (d, *J* = 10.1 Hz, 1H), 2.42 – 2.34 (m, 2H), 2.30 – 2.20 (m, 2H), 2.00 – 1.91 (m, 3H). LC-MS *m/z* calcd. C_30_H_34_N_3_O_2_^+^ [M+H]^+^ = 468.6210, found = 468.3663.

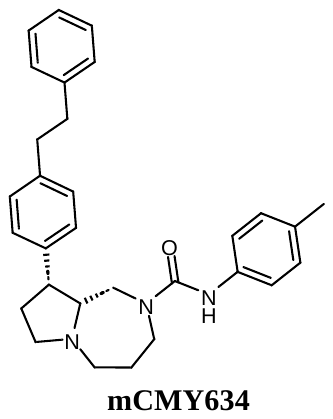

(9R,9aR)-9-(4-phenethylphenyl)-N-(p-tolyl)hexahydro-1H-pyrrolo[1,2-a][1,4]diazepine-2(3H)-carboxamide: ^1^H NMR (400 MHz, Acetone) δ 7.45 (s, 1H), 7.38 (d, *J* = 8.5 Hz, 2H), 7.31 – 7.20 (m, 6H), 7.16 (dd, *J* = 11.2, 7.5 Hz, 3H), 7.01 (d, *J* = 8.2 Hz, 2H), 3.76 (d, *J* = 14.8 Hz, 2H), 3.42 – 3.33 (m, 2H), 3.26 – 3.13 (m, 2H), 2.54 (d, *J* = 5.8 Hz, 2H), 2.42 – 2.28 (m, 3H), 2.23 (d, *J* = 8.5 Hz, 5H), 2.01 – 1.86 (m, 5H). LC-MS *m/z* calcd. C_30_H_36_N_3_O^+^ [M+H]^+^ = 454.42858, found = 454.4629

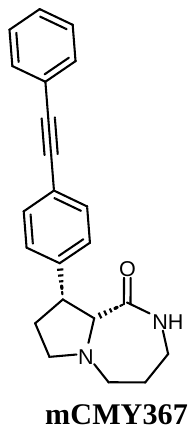

(9R,9aR)-9-(4-(phenylethynyl)phenyl)octahydro-1H-pyrrolo[1,2-a][1,4]diazepin-1-on: ^1^H NMR (400 MHz, Acetone) δ 7.58 – 7.49 (m, 2H), 7.47 – 7.34 (m, 7H), 6.53 (s, 1H), 3.70 – 3.60 (m, 2H), 3.38 (d, *J* = 7.0 Hz, 1H), 3.33 (td, *J* = 8.9, 3.9 Hz, 1H), 3.24 – 3.16 (m, 1H), 2.96 – 2.89 (m, 1H), 2.58 – 2.48 (m, 2H), 2.34 – 2.24 (m, 1H), 2.03 – 1.95 (m, 1H), 1.88 – 1.73 (m, 2H). LC-MS *m/z* calcd. C_22_H_23_N_2_O^+^ [M+H]^+^ = 331.1810, found = 331.2686.

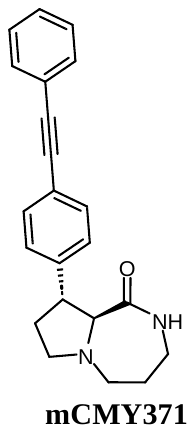

(9R,9aS)-9-(4-(phenylethynyl)phenyl)octahydro-1H-pyrrolo[1,2-a][1,4]diazepin-1-one: ^1^H NMR (400 MHz, Acetone) δ 7.57 – 7.52 (m, 2H), 7.47 (d, *J* = 8.4 Hz, 2H), 7.45 – 7.38 (m, 3H), 7.35 (d, *J* = 8.4 Hz, 2H), 6.88 (s, 1H), 4.26 – 4.16 (m, 1H), 3.37 – 3.20 (m, 5H), 2.83 – 2.76 (m, 1H), 2.58 (dd, *J* = 11.6, 3.9 Hz, 1H), 2.30 – 2.20 (m, 1H), 1.93 – 1.86 (m, 1H), 1.81 – 1.66 (m, 2H). LC-MS *m/z* calcd. C_22_H_23_N_2_O^+^ [M+H]^+^ = 331.1810, found = 331.2686.

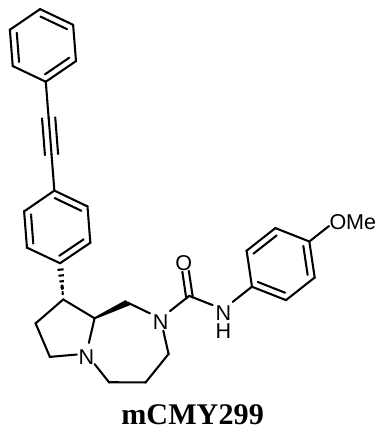

(9R,9aS)-N-(4-methoxyphenyl)-9-(4-(phenylethynyl)phenyl)hexahydro-1H-pyrrolo[1,2-a][1,4]diazepine-2(3H)-carboxamide: ^1^H NMR (400 MHz, CD_3_CN) δ 7.59 – 7.54 (m, 2H), 7.52 (d, *J* = 8.6 Hz, 2H), 7.46 – 7.39 (m, 3H), 7.37 (d, *J* = 7.5 Hz, 2H), 7.22 (d, *J* = 9.0 Hz, 2H), 6.82 (d, *J* = 9.5 Hz, 2H), 6.71 (s, 1H), 3.96 (d, *J* = 13.9 Hz, 1H), 3.73 (s, 3H), 3.58 (t, *J* = 6.6 Hz, 2H), 3.20 – 3.10 (m, 2H), 3.04 – 2.94 (m, 2H), 2.74 – 2.65 (m, 1H), 2.49 (t, *J* = 9.9 Hz, 1H), 2.40 – 2.30 (m, 2H), 1.91 – 1.71 (m, 2H). LC-MS *m/z* calcd. C_30_H_32_N_3_O_2_^+^ [M+H]^+^ = 466.2495, found = 466.3749.

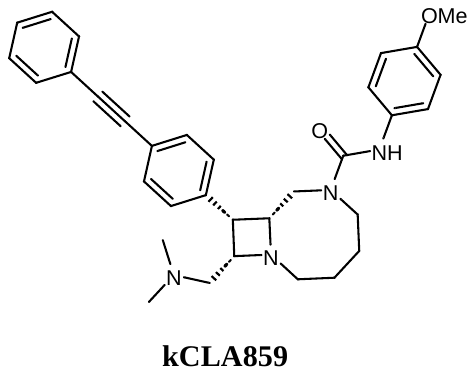

(8R,9S,10S)-10-((dimethylamino)methyl)-N-(4-methoxyphenyl)-9-(4-(phenylethynyl)phenyl)-1,6-diazabicyclo[6.2.0]decane-6-carboxamide: ^1^H NMR (400 MHz, MeOD) δ 7.62 – 7.49 (m, 6H), 7.47 – 7.36 (m, 3H), 7.23 (d, *J* = 8.6 Hz, 2H), 6.87 (d, *J* = 9.2 Hz, 2H), 4.06 (d, *J* = 15.2 Hz, 1H), 3.86 – 3.74 (m, 4H), 3.63 – 3.48 (m, 3H), 3.21 – 2.98 (m, 3H), 2.65 (dd, *J* = 13.4, 8.6 Hz, 1H), 2.56 (d, *J* = 12.6 Hz, 1H), 2.40 – 2.33 (m, 1H), 2.13 (s, 6H), 1.94 – 1.77 (m, 3H), 1.77 – 1.64 (m, 1H). LC-MS *m/z* calcd. C_33_H_39_N_4_O_2_^+^ [M+H]^+^ =523.3073, found = 523.4693.

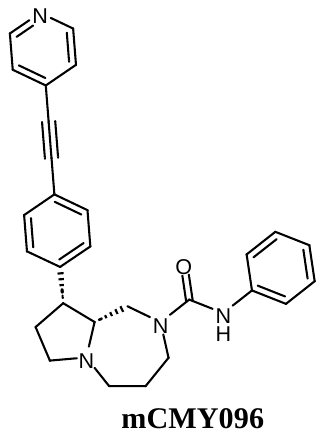

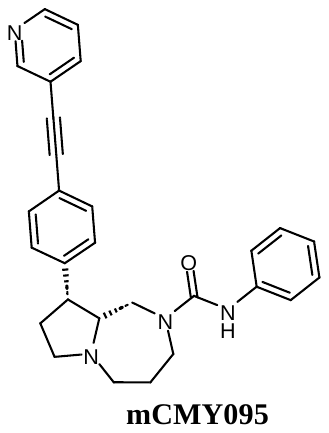
(9R,9aR)-N-phenyl-9-(4-(pyridin-4-ylethynyl)phenyl)hexahydro-1H-pyrrolo[1,2-a][1,4]diazepine-2(3H)-carboxamide: ^1^H NMR (400 MHz, Acetone) δ 8.62 (d, *J* = 6.8 Hz, 2H), 7.59 (s, 1H), 7.55 – 7.46 (m, 8H), 7.21 (d, *J* = 8.2 Hz, 2H), 6.93 (d, *J* = 7.5 Hz, 1H), 3.82 (d, *J* = 14.5 Hz, 1H), 3.75 (dd, *J* = 9.5, 3.9 Hz, 1H), 3.49 – 3.38 (m, 2H), 3.26 (t, *J* = 8.9 Hz, 1H), 3.19 (dt, *J* = 12.6, 4.3 Hz, 1H), 2.60 (t, *J* = 8.3 Hz, 1H), 2.43 – 2.31 (m, 3H), 2.26 – 2.20 (m, 1H), 1.95 (ddd, *J* = 13.1, 7.7, 4.7 Hz, 3H). LC-MS *m/z* calcd. C_28_H_29_N_4_O_2_^+^ [M+H]^+^ =437.2341, found = 437.3722.

(9R,9aR)-N-phenyl-9-(4-(pyridin-3-ylethynyl)phenyl)hexahydro-1H-pyrrolo[1,2-a][1,4]diazepine-2(3H)-carboxamide: ^1^H NMR (400 MHz, CDCl_3_) δ 8.77 (s, 1H), 8.55 (dd, *J* = 4.9, 1.7 Hz, 1H), 7.82 (dt, *J* = 7.9, 1.9 Hz, 1H), 7.53 – 7.47 (m, 2H), 7.47 – 7.39 (m, 2H), 7.39 – 7.33 (m, 2H), 7.32 – 7.28 (m, 3H), 7.04 (t, *J* = 8.5 Hz, 1H), 6.34 (s, 1H), 3.75 (d, *J* = 13.9 Hz, 1H), 3.69 – 3.48 (m, 3H), 3.43 – 3.24 (m, 2H), 2.70 – 2.48 (m, 2H), 2.47 – 2.28 (m, 2H), 2.28 – 2.10 (m, 2H), 2.13 – 1.95 (m, 2H). LC-MS *m/z* calcd. C_28_H_29_N_4_O_2_^+^ [M+H]^+^ =437.2341, found = 437.3730.

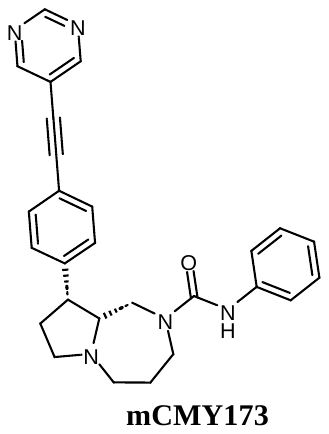

(9R,9aR)-N-phenyl-9-(4-(pyrimidin-5-ylethynyl)phenyl)hexahydro-1H-pyrrolo[1,2-a][1,4]diazepine-2(3H)-carboxamide: ^1^H NMR (400 MHz, Acetone) δ 9.13 (s, 1H), 8.93 (s, 2H), 7.60 (s, 1H), 7.56 – 7.48 (m, 6H), 7.21 (t, *J* = 8.2 Hz, 2H), 6.93 (t, *J* = 7.4 Hz, 1H), 3.82 (d, *J* = 12.9 Hz, 1H), 3.79 – 3.72 (m, 1H), 3.49 – 3.39 (m, 2H), 3.26 (d, *J* = 8.7 Hz, 1H), 3.22 – 3.17 (m, 1H), 2.60 (t, 1H), 2.42 – 2.32 (m, 3H), 2.25 – 2.20 (m, 1H), 1.99 – 1.93 (m, 3H). LC-MS *m/z* calcd. C_27_H_28_N_5_O^+^ [M+H]^+^ =468.2400, found = 438.4229.

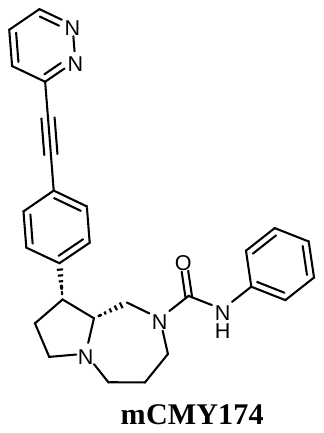

(9R,9aR)-N-phenyl-9-(4-(pyridazin-3-ylethynyl)phenyl)hexahydro-1H-pyrrolo[1,2-a][1,4]diazepine-2(3H)-carboxamide: ^1^H NMR (400 MHz, Acetone) δ 9.19 (dd, *J* = 5.0, 1.7 Hz, 1H), 7.85 (dd, *J* = 8.5, 1.7 Hz, 1H), 7.72 (dd, *J* = 8.5, 5.0 Hz, 1H), 7.61 (d, *J* = 8.4 Hz, 3H), 7.52 (d, *J* = 8.1 Hz, 3H), 7.20 (t, *J* = 8.4 Hz, 2H), 6.93 (t, *J* = 7.5 Hz, 1H), 3.84 (d, *J* = 14.3 Hz, 1H), 3.75 (dd, *J* = 13.5, 6.8 Hz, 1H), 3.52 – 3.45 (m, 1H), 3.45 – 3.37 (m, 1H), 3.26 (t, *J* = 8.4 Hz, 1H), 3.19 (td, *J* = 13.0, 3.6 Hz, 1H), 2.60 (t, *J* = 8.0 Hz, 1H), 2.44 – 2.32 (m, 3H), 2.28 – 2.20 (m, 1H), 2.01 – 1.91 (m, 3H). LC-MS *m/z* calcd. C_27_H_28_N_5_O^+^ [M+H]^+^ =468.2400, found = 438.3992.

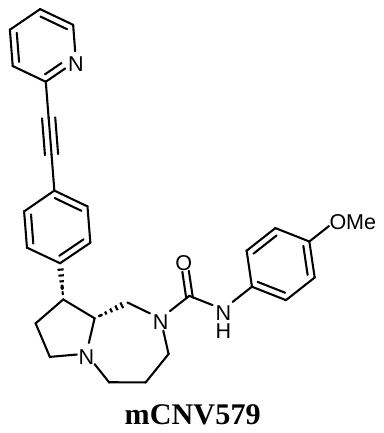

(9R,9aR)-N-(4-methoxyphenyl)-9-(4-(pyridin-2-ylethynyl)phenyl)hexahydro-1H-pyrrolo[1,2-a][1,4]diazepine-2(3H)-carboxamide: ^1^H NMR (400 MHz, DMSO) δ 8.62 (d, *J* = 4.9 Hz, 1H), 8.00 (s, 1H), 7.86 (t, *J* = 7.7 Hz, 1H), 7.64 (d, *J* = 7.9 Hz, 1H), 7.55 (d, *J* = 7.6 Hz, 2H), 7.43 (d, *J* = 7.6 Hz, 3H), 7.31 (d, *J* = 8.3 Hz, 2H), 6.80 (d, *J* = 8.3 Hz, 2H), 3.70 (d, *J* = 1.5 Hz, 5H), 3.24 – 3.07 (m, 4H), 2.39 (s, 4H), 2.31 – 2.09 (m, 2H), 1.87 (s, 3H). LC-MS *m/z* calcd. C_29_H_30_N_4_O_2_^+^ [M+H]^+^ = 467.2369, found = 467.4611.

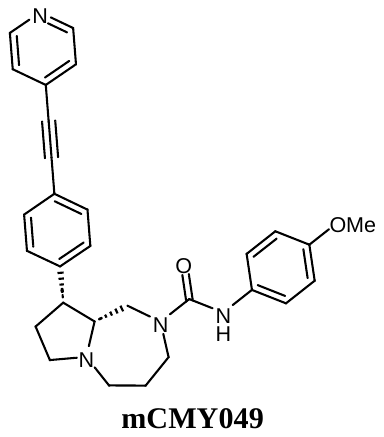

(9R,9aR)-N-(4-methoxyphenyl)-9-(4-(pyridin-4-ylethynyl)phenyl)hexahydro-1H-pyrrolo[1,2-a][1,4]diazepine-2(3H)-carboxamide: ^1^H NMR (400 MHz, CDCl_3_) δ 8.62 (d, *J* = 5.2 Hz, 2H), 7.50 (d, *J* = 8.2 Hz, 2H), 7.46 – 7.32 (m, 4H), 7.26 (d, *J* = 8.1 Hz, 2H), 6.84 (d, 2H), 6.20 (s, 1H), 3.79 (s, 4H), 3.55 (t, *J* = 6.3 Hz, 2H), 3.52 – 3.43 (m, 1H), 3.37 – 3.20 (m, 2H), 2.81 – 2.63 (m, 1H), 2.59 – 2.40 (m, 2H), 2.40 – 2.25 (m, 2H), 2.13 – 1.95 (m, 3H). LC-MS *m/z* calcd. C_29_H_30_N_4_O_2_^+^ [M+H]^+^ = 467.2447, found = 467.4249.

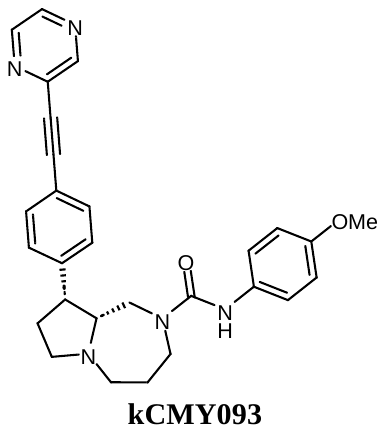

(9R,9aR)-N-(4-methoxyphenyl)-9-(4-(pyrazin-2-ylethynyl)phenyl)hexahydro-1H-pyrrolo[1,2-a][1,4]diazepine-2(3H)-carboxamide: ^1^H NMR (400 MHz, MeOD) δ 8.81 (s, 1H), 8.62 (dd, *J* = 2.6, 1.6 Hz, 1H), 8.56 (d, *J* = 2.6 Hz, 1H), 7.58 (d, *J* = 8.1 Hz, 2H), 7.48 (d, *J* = 8.4 Hz, 2H), 7.19 (d, *J* = 8.8 Hz, 2H), 6.83 (d, *J* = 9.1 Hz, 2H), 3.76 (s, 3H), 3.71 – 3.60 (m, 2H), 3.58 – 3.53 (m, 1H), 3.44 (dt, *J* = 13.6, 5.6 Hz, 1H), 3.29 – 3.21 (m, 2H), 2.70 (t, *J* = 9.8 Hz, 1H), 2.53 – 2.45 (m, 2H), 2.35 – 2.26 (m, 2H), 2.08 – 1.97 (m, 3H). LC-MS *m/z* calcd. C_28_H_29_N_5_O_2_^+^ [M+H]^+^ = 468.2400, found = 468.4341.

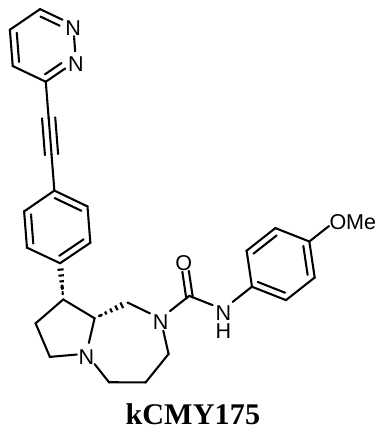

(9R,9aR)-N-(4-methoxyphenyl)-9-(4-(pyridazin-3-ylethynyl)phenyl)hexahydro-1H-pyrrolo[1,2-a][1,4]diazepine-2(3H)-carboxamide: ^1^H NMR (400 MHz, Acetone) δ 9.19 (d, *J* = 5.0 Hz, 1H), 7.85 (dd, *J* = 8.5, 1.7 Hz, 1H), 7.72 (dd, *J* = 8.5, 5.0 Hz, 1H), 7.61 (d, *J* = 8.1 Hz, 2H), 7.51 (d, *J* = 8.1 Hz, 2H), 7.45 (s, 1H), 7.39 (d, *J* = 9.1 Hz, 2H), 6.79 (d, *J* = 9.1 Hz, 2H), 3.83 (d, *J* = 13.1 Hz, 1H), 3.74 (s, 3H), 3.48 (d, *J* = 5.5 Hz, 1H), 3.39 (d, *J* = 13.2 Hz, 1H), 3.29 – 3.17 (m, 2H), 2.63 – 2.59 (m, 1H), 2.45 – 2.33 (m, 3H), 2.24 – 2.20 (m, 1H), 2.00 – 1.88 (m, 4H). LC-MS *m/z* calcd. C_28_H_29_N_5_O_2_^+^ [M+H]^+^ = 468.2400, found = 468.4335.

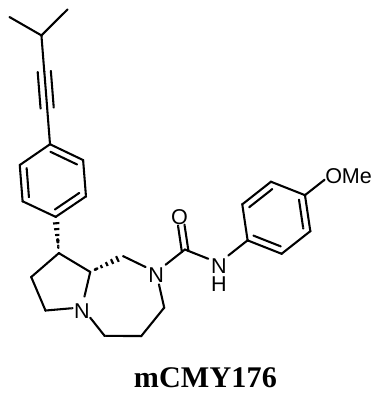

(9R,9aR)-N-(4-methoxyphenyl)-9-(4-(3-methylbut-1-yn-1-yl)phenyl)hexahydro-1H-pyrrolo[1,2-a][1,4]diazepine-2(3H)-carboxamide: ^1^H NMR (400 MHz, Acetone) δ 7.43 (s, 1H), 7.38 (d, *J* = 9.1 Hz, 2H), 7.36 – 7.27 (m, 4H), 6.79 (d, *J* = 8.7 Hz, 2H), 3.77 (d, *J* = 12.9 Hz, 1H), 3.74 (s, 3H), 3.42 – 3.32 (m, 2H), 3.22 (d, *J* = 8.9 Hz, 1H), 3.19 – 3.14 (m, 1H), 2.81 – 2.75 (m, 1H), 2.56 (d, *J* = 8.9 Hz, 1H), 2.42 – 2.31 (m, 2H), 2.32 – 2.15 (m, 3H), 1.98 – 1.87 (m, 3H), 1.24 (d, *J* = 6.8 Hz, 6H). LC-MS *m/z* calcd. C_27_H_34_N_3_O_2_^+^ [M+H]^+^ = 432.2651, found = 432.4846.

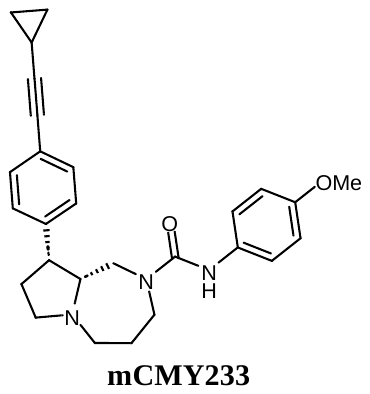

(9R,9aR)-9-(4-(cyclopropylethynyl)phenyl)-N-(4-methoxyphenyl)hexahydro-1H-pyrrolo[1,2-a][1,4]diazepine-2(3H)-carboxamide: ^1^H NMR (400 MHz, Acetone) δ 7.45 (s, 1H), 7.38 (d, *J* = 9.3 Hz, 2H), 7.35 – 7.24 (m, 4H), 6.79 (d, *J* = 8.8 Hz, 2H), 3.79 – 3.68 (m, 5H), 3.40 – 3.33 (m, 2H), 3.21 (t, *J* = 8.5 Hz, 1H), 3.16 (dt, *J* = 12.6, 4.3 Hz, 1H), 2.55 (t, *J* = 8.7 Hz, 1H), 2.42 – 2.30 (m, 2H), 2.28 – 2.15 (m, 2H), 1.97 – 1.86 (m, 3H), 1.54 – 1.44 (m, 1H), 0.92 – 0.86 (m, 2H), 0.76 – 0.65 (m, 2H). LC-MS *m/z* calcd. C_27_H_32_N_3_O_2_^+^ [M+H]^+^ = 430.2495, found = 430.4569.

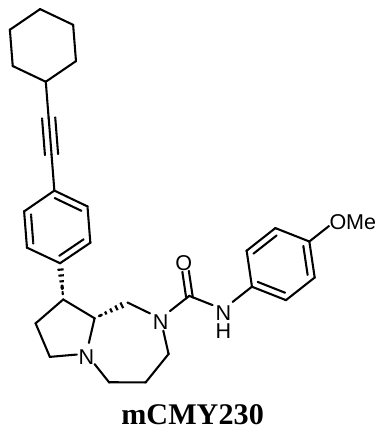

(9R,9aR)-9-(4-(cyclohexylethynyl)phenyl)-N-(4-methoxyphenyl)hexahydro-1H-pyrrolo[1,2-a][1,4]diazepine-2(3H)-carboxamide: ^1^H NMR (400 MHz, Acetone) δ 7.45 (s, 1H), 7.38 (d, *J* = 9.1 Hz, 2H), 7.35 – 7.29 (m, 4H), 6.79 (d, *J* = 9.1 Hz, 2H), 3.80 – 3.69 (m, 5H), 3.41 – 3.34 (m, 2H), 3.22 (d, *J* = 8.0 Hz, 1H), 3.15 (dt, *J* = 12.6, 3.8 Hz, 1H), 2.66 – 2.59 (m, 1H), 2.55 (t, *J* = 8.2 Hz, 1H), 2.40 – 2.34 (m, 1H), 2.31 – 2.26 (m, 1H), 2.24 – 2.17 (m, 1H), 1.97 – 1.84 (m, 5H), 1.79 – 1.72 (m, 2H), 1.58 – 1.47 (m, 3H), 1.43 – 1.35 (m, 3H). LC-MS *m/z* calcd. C_30_H_38_N_3_O_2_^+^ [M+H]^+^ = 472.2964, found = 472.5300.

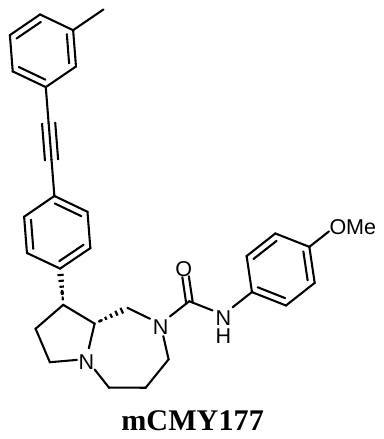

(9R,9aR)-N-(4-methoxyphenyl)-9-(4-(m-tolylethynyl)phenyl)hexahydro-1H-pyrrolo[1,2-a][1,4]diazepine-2(3H)-carboxamide: ^1^H NMR (400 MHz, Acetone) δ 7.49 – 7.42 (m, 5H), 7.40 – 7.33 (m, 4H), 7.30 (d, *J* = 6.9 Hz, 1H), 7.22 (d, *J* = 7.4 Hz, 1H), 6.79 (d, *J* = 8.8 Hz, 2H), 3.81 (d, 1H), 3.74 (s, 4H), 3.47 – 3.34 (m, 2H), 3.25 (t, *J* = 8.6 Hz, 1H), 3.18 (dt, *J* = 12.6, 4.2 Hz, 1H), 2.58 (t, *J* = 8.8 Hz, 1H), 2.44 – 2.38 (m, 1H), 2.35 (s, 4H), 2.34 – 2.28 (m, 1H), 2.28 – 2.18 (m, 1H), 1.98 – 1.89 (m, 3H). LC-MS *m/z* calcd. C_31_H_34_N_3_O_2_^+^ [M+H]^+^ = 480.2651, found = 480.4952

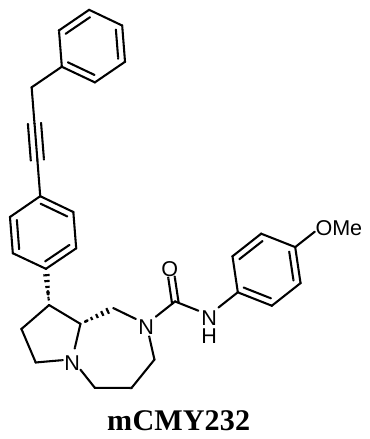

(9R,9aR)-N-(4-methoxyphenyl)-9-(4-(3-phenylprop-1-yn-1-yl)phenyl)hexahydro-1H-pyrrolo[1,2-a][1,4]diazepine-2(3H)-carboxamide: ^1^H NMR (400 MHz, Acetone) δ 7.45 (d, *J* = 7.8 Hz, 3H), 7.43 – 7.31 (m, 8H), 7.26 (t, *J* = 6.8 Hz, 1H), 6.79 (d, *J* = 9.3 Hz, 2H), 3.88 (s, 2H), 3.81 – 3.70 (m, 5H), 3.41 (s, 2H), 3.29 – 3.15 (m, 2H), 2.68 – 2.53 (m, 1H), 2.48 – 2.21 (m, 4H), 2.02 – 1.88 (m, 3H). LC-MS *m/z* calcd. C_31_H_34_N_3_O_2_^+^ [M+H]^+^ = 480.2651, found = 480.4558.

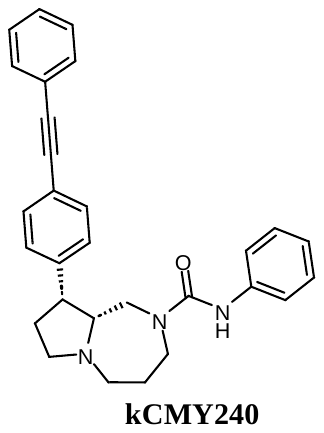

(9R,9aR)-N-phenyl-9-(4-(phenylethynyl)phenyl)hexahydro-1H-pyrrolo[1,2-a][1,4]diazepine-2(3H)-carboxamide: ^1^H NMR (400 MHz, Acetone) δ 7.63 – 7.53 (m, 3H), 7.53 – 7.38 (m, 8H), 7.21 (t, *J* = 7.5 Hz, 2H), 6.92 (tt, *J* = 7.5, 1.1 Hz, 1H), 3.82 (d, *J* = 14.5 Hz, 1H), 3.79 – 3.70 (m, 1H), 3.49 – 3.38 (m, 2H), 3.31 – 3.14 (m, 2H), 2.64 – 2.57 (m, 1H), 2.47 – 2.28 (m, 3H), 2.27 – 2.20 (m, 1H), 2.02 – 1.84 (m, 3H). LC-MS *m/z* calcd. C_29_H_30_N_3_O^+^ [M+H]^+^ = 436.2389, found = 436.4008.

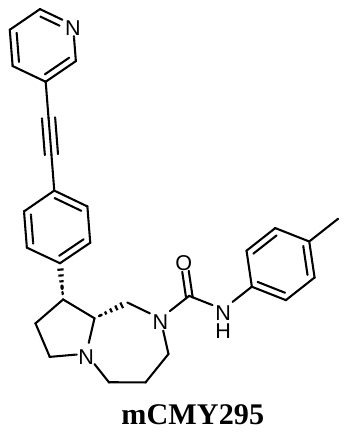

(9R,9aR)-9-(4-(pyridin-3-ylethynyl)phenyl)-N-(p-tolyl)hexahydro-1H-pyrrolo[1,2-a][1,4]diazepine-2(3H)-carboxamide: ^1^H NMR (400 MHz, Acetone) δ 8.74 (s, 1H), 8.57 (dd, *J* = 4.9, 1.7 Hz, 1H), 7.92 (dt, *J* = 7.9, 1.7 Hz, 1H), 7.55 – 7.40 (m, 6H), 7.38 (d, *J* = 9.0 Hz, 2H), 7.02 (d, *J* = 8.4 Hz, 2H), 3.82 (d, *J* = 14.3 Hz, 1H), 3.73 (dd, *J* = 13.9, 6.0 Hz, 1H), 3.48 – 3.36 (m, 2H), 3.25 (t, *J* = 8.8 Hz, 1H), 3.18 (dt, *J* = 13.1, 4.0 Hz, 1H), 2.62 – 2.57 (m, 1H), 2.43 – 2.31 (m, 3H), 2.26 – 2.20 (m, 4H), 1.99 – 1.90 (m, 3H). LC-MS *m/z* calcd. C_29_H_31_N_4_O^+^ [M+H]^+^ = 451.2498, found = 451.3491.

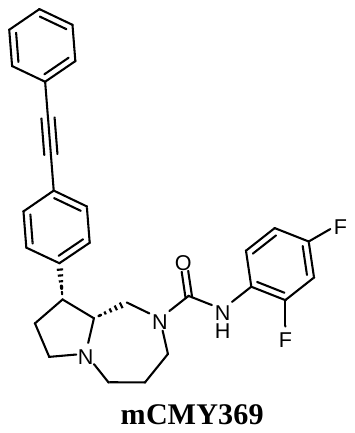

(9R,9aR)-N-(2,4-difluorophenyl)-9-(4-(phenylethynyl)phenyl)hexahydro-1H-pyrrolo[1,2-a][1,4]diazepine-2(3H)-carboxamide: ^1^H NMR (400 MHz, Acetone) δ 7.81 – 7.71 (m, 1H), 7.58 – 7.52 (m, 2H), 7.49 (d, *J* = 8.4 Hz, 2H), 7.46 – 7.37 (m, 5H), 7.23 (s, 1H), 7.01 (t, *J* = 10.8 Hz, 1H), 6.91 (t, *J* = 9.0 Hz, 1H), 3.82 – 3.68 (m, 2H), 3.51 – 3.40 (m, 2H), 3.30 – 3.13 (m, 2H), 2.62 (t, *J* = 8.9 Hz, 1H), 2.49 – 2.37 (m, 2H), 2.35 – 2.21 (m, 2H), 2.01 – 1.91 (m, 3H). LC-MS *m/z* calcd. C_29_H_28_F_2_N_3_O^+^ [M+H]^+^ = 472.2200, found = 472.3851.

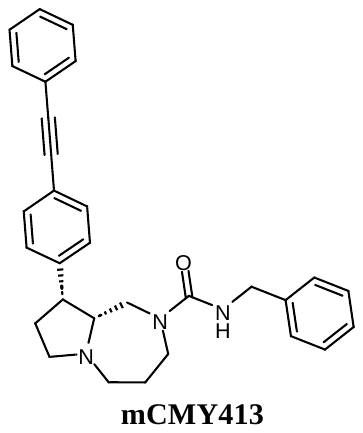

(9R,9aR)-N-benzyl-9-(4-(phenylethynyl)phenyl)hexahydro-1H-pyrrolo[1,2-a][1,4]diazepine-2(3H)-carboxamide: ^1^H NMR (400 MHz, Acetone) δ 7.58 – 7.53 (m, 2H), 7.46 (d, J = 8.4 Hz, 2H), 7.44 – 7.37 (m, 5H), 7.30 – 7.24 (m, 4H), 7.22 – 7.16 (m, 1H), 4.34 (d, J = 5.9 Hz, 2H), 3.75 (d, J = 14.3 Hz, 1H), 3.71 – 3.59 (m, 1H), 3.44 – 3.35 (m, 1H), 3.32 – 3.20 (m, 2H), 3.15 (dt, J = 12.8, 4.0 Hz, 1H), 2.52 (t, J = 8.7 Hz, 1H), 2.41 – 2.25 (m, 3H), 2.19 (t, J = 11.3 Hz, 1H), 2.10 (s, 2H), 1.97 – 1.84 (m, 2H). LC-MS *m/z* calcd. C_30_H_32_N_3_O^+^ [M+H]^+^ = 450.2545, found = 450.4077.

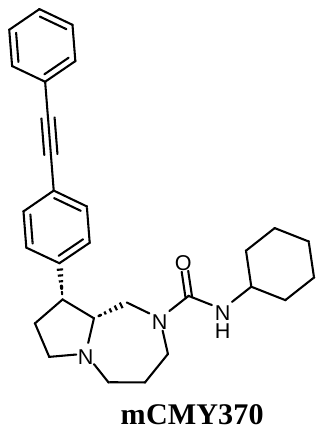

(9R,9aR)-N-cyclohexyl-9-(4-(phenylethynyl)phenyl)hexahydro-1H-pyrrolo[1,2-a][1,4]diazepine-2(3H)-carboxamide: ^1^H NMR (400 MHz, Acetone) δ 7.56 – 7.53 (m, 2H), 7.47 (d, *J* = 8.3 Hz, 2H), 7.44 – 7.39 (m, 5H), 5.16 (d, *J* = 7.8 Hz, 1H), 3.74 (d, *J* = 14.6 Hz, 1H), 3.58 – 3.51 (m, 2H), 3.42 – 3.36 (m, 1H), 3.25 – 3.18 (m, 2H), 3.13 (dt, *J* = 12.4, 4.1 Hz, 1H), 2.49 (t, *J* = 8.4 Hz, 1H), 2.39 – 2.34 (m, 1H), 2.24 – 2.13 (m, 2H), 1.93 – 1.82 (m, 5H), 1.71 – 1.66 (m, 2H), 1.61 – 1.55 (m, 1H), 1.32 – 1.26 (m, 3H), 1.18 – 1.10 (m, 3H). LC-MS *m/z* calcd. C_29_H_36_N_3_O^+^ [M+H]^+^ = 442.2858, found = 442.4058.

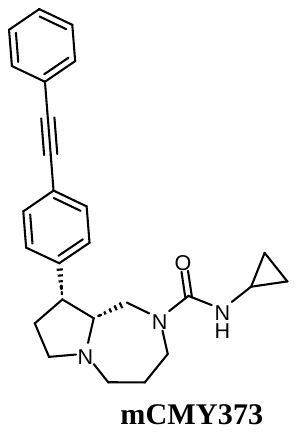

(9R,9aR)-N-cyclopropyl-9-(4-(phenylethynyl)phenyl)hexahydro-1H-pyrrolo[1,2-a][1,4]diazepine-2(3H)-carboxamide: ^1^H NMR (400 MHz, Acetone) δ 7.58 – 7.51 (m, 2H), 7.46 (d, *J* = 8.4 Hz, 2H), 7.44 – 7.37 (m, 5H), 5.69 (s, 1H), 3.70 (d, *J* = 14.3 Hz, 1H), 3.57 – 3.47 (m, 1H), 3.43 – 3.35 (m, 1H), 3.24 – 3.10 (m, 3H), 2.60 – 2.52 (m, 1H), 2.48 (t, *J* = 9.8 Hz, 1H), 2.40 – 2.13 (m, 4H), 1.95 – 1.81 (m, 3H), 0.59 – 0.49 (m, 2H), 0.39 – 0.32 (m, 2H). LC-MS *m/z* calcd. C_26_H_30_N_3_O^+^ [M+H]^+^ = 400.2389, found = 400.4387.

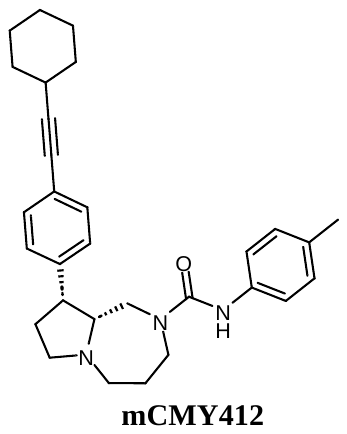

(9R,9aR)-9-(4-(cyclohexylethynyl)phenyl)-N-(p-tolyl)hexahydro-1H-pyrrolo[1,2-a][1,4]diazepine-2(3H)-carboxamide: ^1^H NMR (400 MHz, Acetone) δ 7.48 (s, 1H), 7.41 – 7.26 (m, 6H), 7.01 (d, *J* = 8.3 Hz, 2H), 3.81 – 3.68 (m, 2H), 3.44 – 3.34 (m, 2H), 3.25 – 3.13 (m, 2H), 2.64 – 2.53 (m, 2H), 2.42 – 2.27 (m, 3H), 2.25 – 2.18 (m, 4H), 1.97 – 1.85 (m, 5H), 1.79 – 1.72 (m, 2H), 1.59 – 1.48 (m, 3H), 1.42 – 1.34 (m, 3H). LC-MS *m/z* calcd. C_30_H_38_N_3_O^+^ [M+H]^+^ = 456.3015, found = 456.8526.

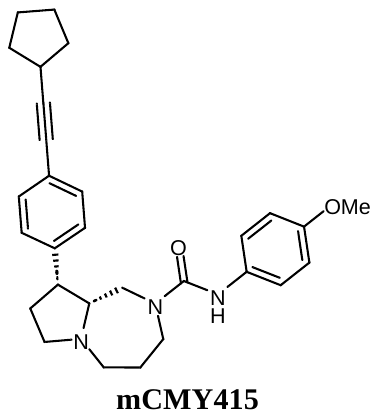

(9R,9aR)-9-(4-(cyclopentylethynyl)phenyl)-N-(4-methoxyphenyl)hexahydro-1H-pyrrolo[1,2-a][1,4]diazepine-2(3H)-carboxamide: ^1^H NMR (400 MHz, Acetone) δ 7.54 (s, 1H), 7.38 – 7.32 (m, 4H), 7.28 (d, *J* = 8.6 Hz, 2H), 6.78 (d, *J* = 8.6 Hz, 2H), 3.78 – 3.67 (m, 5H), 3.45 – 3.33 (m, 3H), 2.89 – 2.78 (m, 1H), 2.61 – 2.52 (m, 1H), 2.40 – 2.15 (m, 4H), 2.03 – 1.86 (m, 6Y415H), 1.81 – 1.71 (m, 2H), 1.71 – 1.56 (m, 4H). LC-MS *m/z* calcd. C_29_H_36_N_3_O_2_^+^ [M+H]^+^ = 458.2808, found = 458.4819.

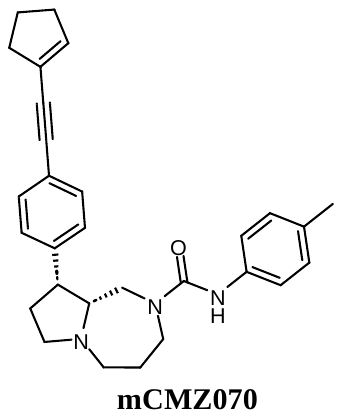

(9R,9aR)-9-(4-(cyclopent-1-en-1-ylethynyl)phenyl)-N-(p-tolyl)hexahydro-1H-pyrrolo[1,2-a][1,4]diazepine-2(3H)-carboxamide: ^1^H NMR (400 MHz, Acetone) δ 7.49 (s, 1H), 7.42 – 7.31 (m, 6H), 7.01 (d, *J* = 8.1 Hz, 2H), 6.10 (ddd, *J* = 4.7, 2.7, 1.9 Hz, 1H), 3.79 (d, *J* = 15.3 Hz, 1H), 3.76 – 3.66 (m, 1H), 3.44 – 3.35 (m, 2H), 3.23 (td, *J* = 8.5, 1.9 Hz, 1H), 3.17 (dt, *J* = 12.6, 4.2 Hz, 1H), 2.60 – 2.54 (m, 1H), 2.54 – 2.45 (m, 4H), 2.43 – 2.33 (m, 2H), 2.33 – 2.29 (m, 1H), 2.24 (s, 3H), 2.23 – 2.17 (m, 1H), 1.99 – 1.89 (m, 5H). LC-MS *m/z* calcd. C_29_H_34_N_3_O_2_^+^ [M+H]^+^ = 440.2702, found = 440.4143.

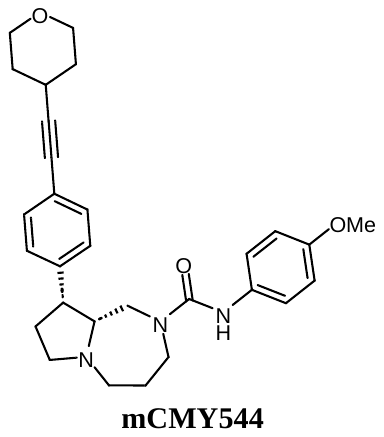

(9R,9aR)-N-(4-methoxyphenyl)-9-(4-((tetrahydro-2H-pyran-4-yl)ethynyl)phenyl)hexahydro-1H-pyrrolo[1,2-a][1,4]diazepine-2(3H)-carboxamide: ^1^H NMR (400 MHz, Acetone) δ 7.48 (s, 1H), 7.45 (d, *J* = 8.3 Hz, 1H), 7.38 (d, *J* = 8.4 Hz, 2H), 7.35 – 7.31 (m, 3H), 6.79 (d, *J* = 9.2 Hz, 2H), 3.91 – 3.80 (m, 2H), 3.80 – 3.67 (m, 5H), 3.49 (t, *J* = 10.7 Hz, 1H), 3.45 – 3.30 (m, 3H), 3.24 – 3.14 (m, 2H), 2.90 – 2.81 (m, 1H), 2.58 – 2.51 (m, 1H), 2.41 – 2.27 (m, 3H), 2.24 – 2.16 (m, 1H), 2.03 – 1.79 (m, 5H), 1.72 – 1.54 (m, 2H). LC-MS *m/z* calcd. C_29_H_36_N_3_O_3_^+^ [M+H]^+^ = 474.2757, found = 474.4851.

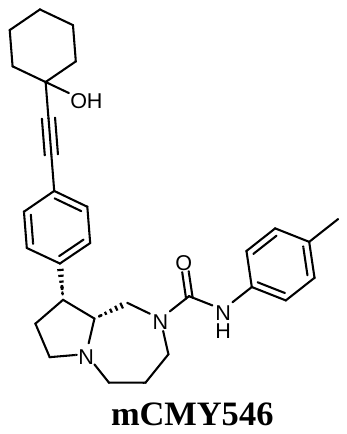

(9R,9aR)-9-(4-((1-hydroxycyclohexyl)ethynyl)phenyl)-N-(p-tolyl)hexahydro-1H-pyrrolo[1,2-a][1,4]diazepine-2(3H)-carboxamide: ^1^H NMR (400 MHz, Acetone) δ 7.49 (s, 1H), 7.44 – 7.30 (m, 6H), 7.03 (d, *J* = 8.2 Hz, 2H), 4.42 (s, 1H), 3.83 – 3.70 (m, 2H), 3.47 – 3.37 (m, 2H), 3.27 – 3.16 (m, 2H), 2.58 (t, *J* = 8.7 Hz, 1H), 2.45 – 2.31 (m, 3H), 2.27 – 2.22 (m, 4H), 1.99 – 1.89 (m, 5H), 1.77 – 1.53 (m, 7H), 1.37 – 1.28 (m, 1H). LC-MS *m/z* calcd. C_30_H_38_N_3_O_2_^+^ [M+H]^+^ = 472.2964, found = 472.4214.

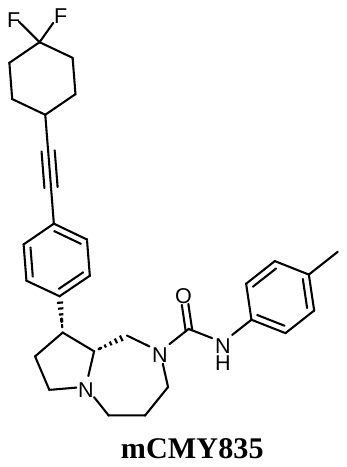

(9R,9aR)-9-(4-((4,4-difluorocyclohexyl)ethynyl)phenyl)-N-(p-tolyl)hexahydro-1H-pyrrolo[1,2-a][1,4]diazepine-2(3H)-carboxamide: ^1^H NMR (400 MHz, Acetone) δ 7.48 (s, 1H), 7.39 – 7.32 (m, 6H), 7.01 (d, *J* = 8.6 Hz, 2H), 3.78 (d, *J* = 13.1 Hz, 1H), 3.76 – 3.68 (m, 1H), 3.43 – 3.34 (m, 2H), 3.22 (td, *J* = 8.6, 2.0 Hz, 1H), 3.18 – 3.13 (m, 1H), 2.54 (t, *J* = 9.2 Hz, 1H), 2.42 – 2.33 (m, 2H), 2.32 – 2.26 (m, 2H), 2.24 (s, 3H), 2.22 – 2.12 (m, 3H), 2.01 – 1.90 (m, 7H), 1.84 – 1.77 (m, 2H). LC-MS *m/z* calcd. C_30_H_36_F_2_N_3_O^+^ [M+H]^+^ = 492.2826, found = 492.4428.

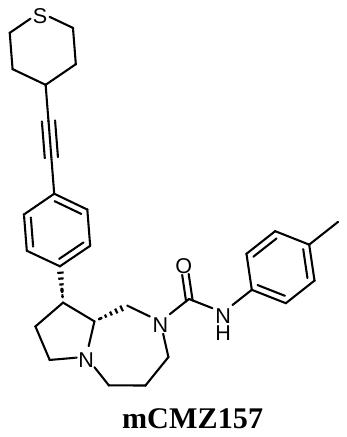

(9R,9aR)-9-(4-((tetrahydro-2H-thiopyran-4-yl)ethynyl)phenyl)-N-(p-tolyl)hexahydro-1H-pyrrolo[1,2-a][1,4]diazepine-2(3H)-carboxamide: ^1^H NMR (400 MHz, Acetone) δ 7.47 (s, 1H), 7.40 – 7.31 (m, 6H), 7.01 (d, *J* = 8.7 Hz, 2H), 3.82 – 3.69 (m, 2H), 3.45 – 3.34 (m, 2H), 3.27 – 3.12 (m, 2H), 2.63 – 2.54 (m, 3H), 2.43 – 2.33 (m, 2H), 2.33 – 2.24 (m, 2H), 2.24 (s, 3H), 2.23 – 2.20 (m, 1H), 2.20 – 2.11 (m, 2H), 2.02 – 1.84 (m, 7H). LC-MS *m/z* calcd. C_29_H_36_N_3_OS^+^ [M+H]^+^ = 474.2579, found = 474.3765.

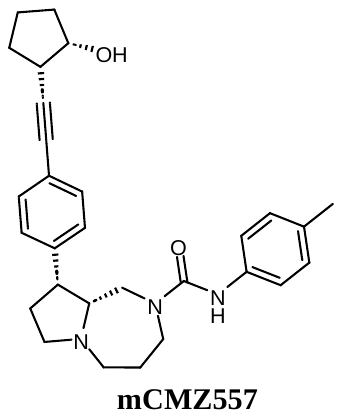

(9R,9aR)-9-(4-(((1S,2S)-2-hydroxycyclopentyl)ethynyl)phenyl)-N-(p-tolyl)hexahydro-1H-pyrrolo[1,2-a][1,4]diazepine-2(3H)-carboxamide^: 1^H NMR (400 MHz, Acetone) δ 7.48 (s, 1H), 7.41 – 7.34 (m, 4H), 7.31 (d, *J* = 8.0 Hz, 2H), 7.03 (d, *J* = 8.3 Hz, 2H), 4.28 – 4.23 (m, 1H), 4.12 (d, *J* = 4.2 Hz, 1H), 3.82 – 3.70 (m, 2H), 3.44 – 3.36 (m, 2H), 3.26 – 3.16 (m, 2H), 2.81 – 2.76 (m, 1H), 2.60 – 2.55 (m, 1H), 2.42 – 2.31 (m, 3H), 2.26 – 2.20 (m, 5H), 2.01 – 1.91 (m, 4H), 1.81 – 1.70 (m, 3H), 1.65 – 1.59 (m, 1H). LC-MS *m/z* calcd. C_29_H_36_N_3_O_2_^+^ [M+H]^+^ = 458.2808, found = 458.3733.

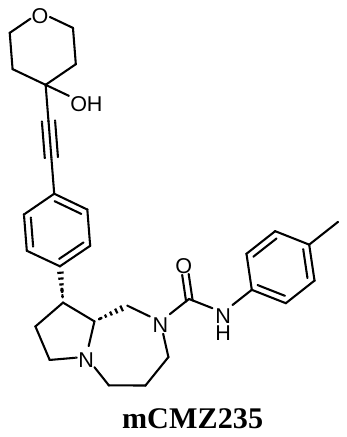

(9R,9aR)-9-(4-((4-hydroxytetrahydro-2H-pyran-4-yl)ethynyl)phenyl)-N-(p-tolyl)hexahydro-1H-pyrrolo[1,2-a][1,4]diazepine-2(3H)-carboxamide: ^1^H NMR (400 MHz, Acetone) δ 7.48 (s, 1H), 7.42 – 7.33 (m, 6H), 7.01 (d, *J* = 8.4 Hz, 2H), 4.72 (s, 1H), 3.90 – 3.82 (m, 2H), 3.81 – 3.76 (m, 1H), 3.75 – 3.69 (m, 1H), 3.65 (ddd, *J* = 11.6, 8.8, 3.0 Hz, 2H), 3.45 – 3.35 (m, 2H), 3.23 (t, *J* = 9.3 Hz, 1H), 3.17 (dt, *J* = 12.8, 4.9 Hz, 1H), 2.57 (t, *J* = 9.6 Hz, 1H), 2.43 – 2.28 (m, 3H), 2.24 (s, 3H), 2.23 – 2.18 (m, 1H), 1.98 – 1.89 (m, 5H), 1.80 (ddd, *J* = 12.9, 8.9, 3.9 Hz, 2H). LC-MS *m/z* calcd. C_29_H_36_N_3_O_3_^+^ [M+H]^+^ = 474.2757, found = 474.3402.

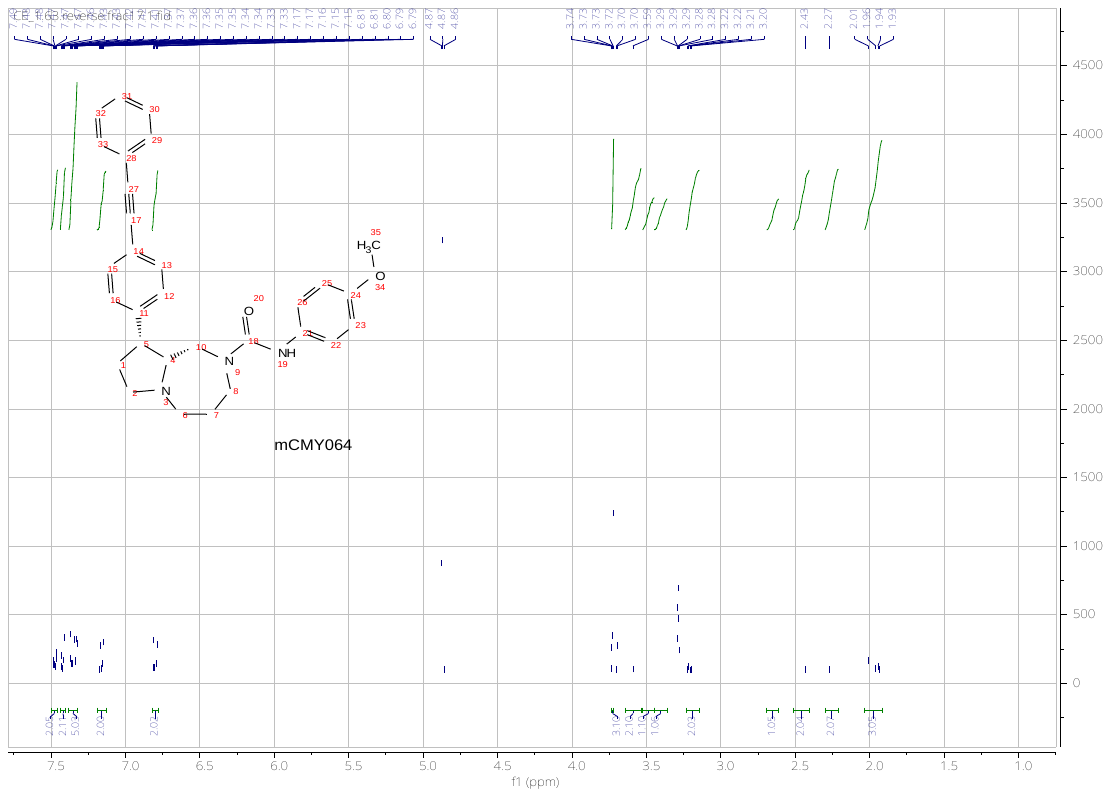

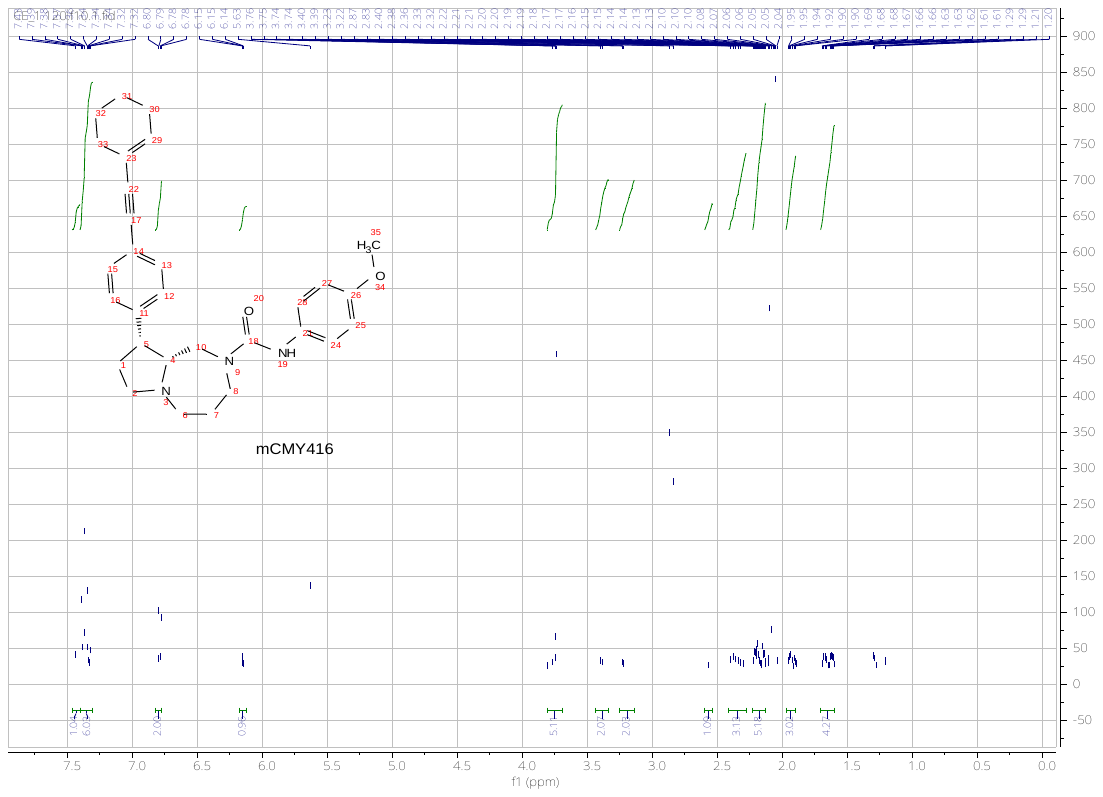

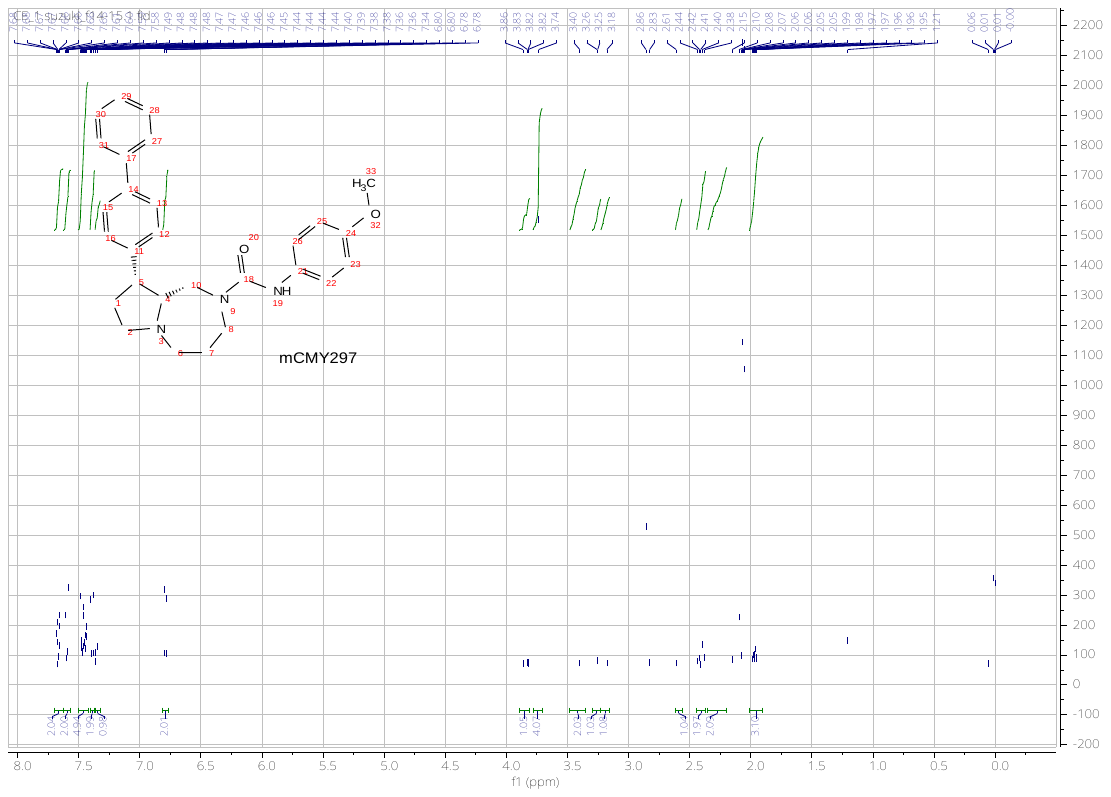

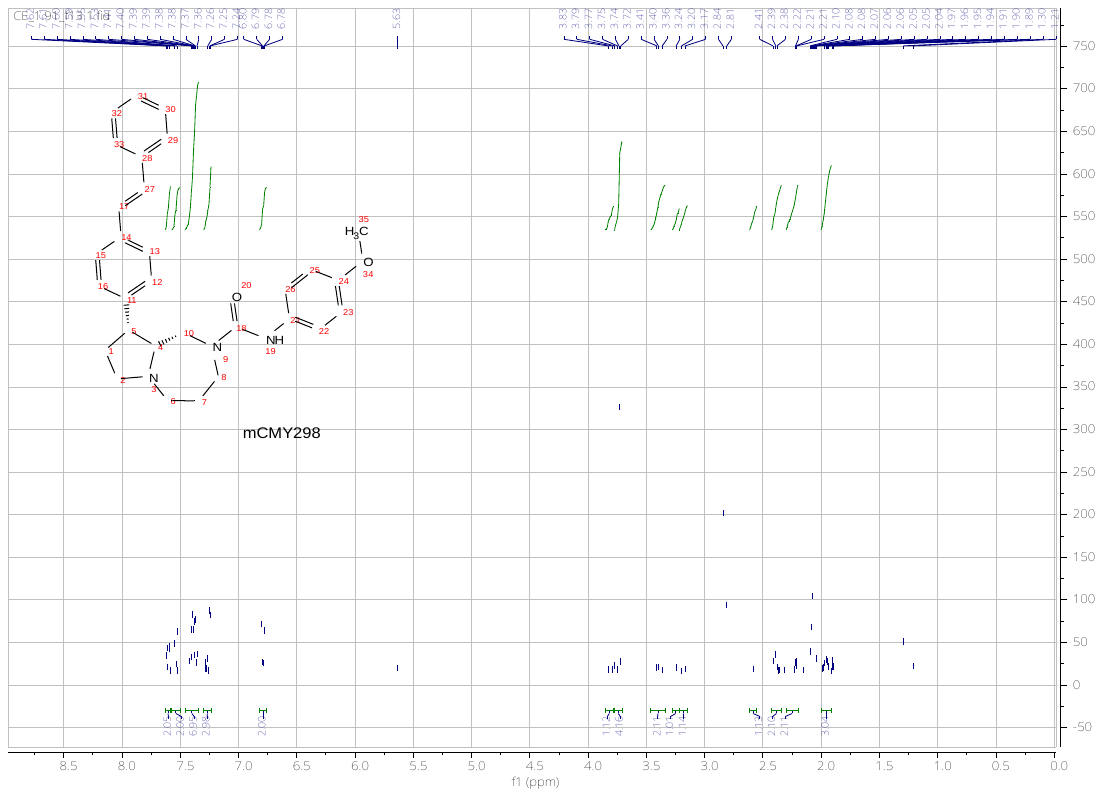

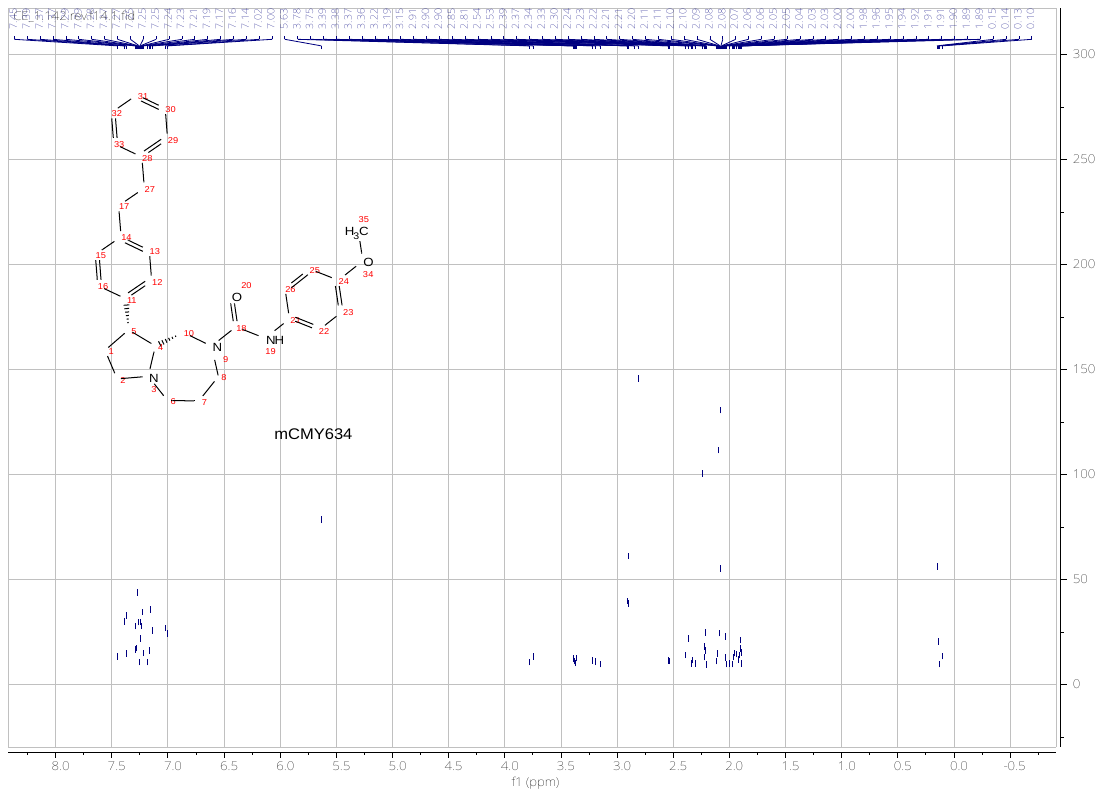

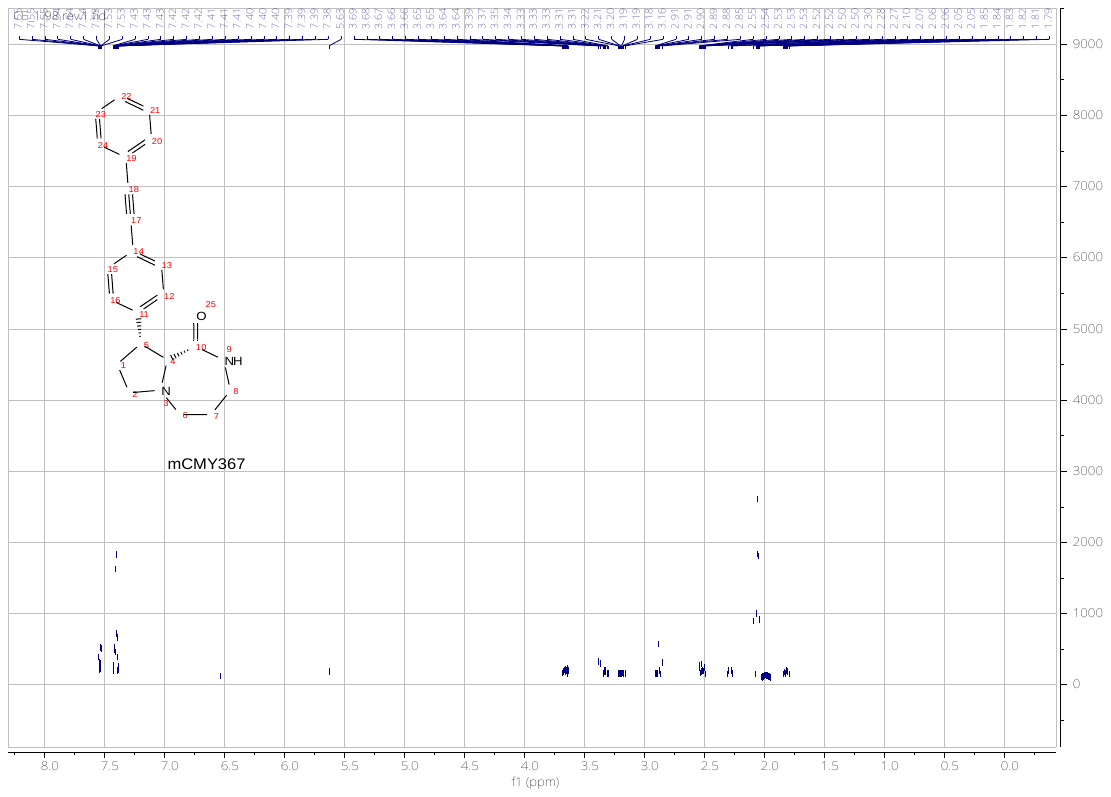

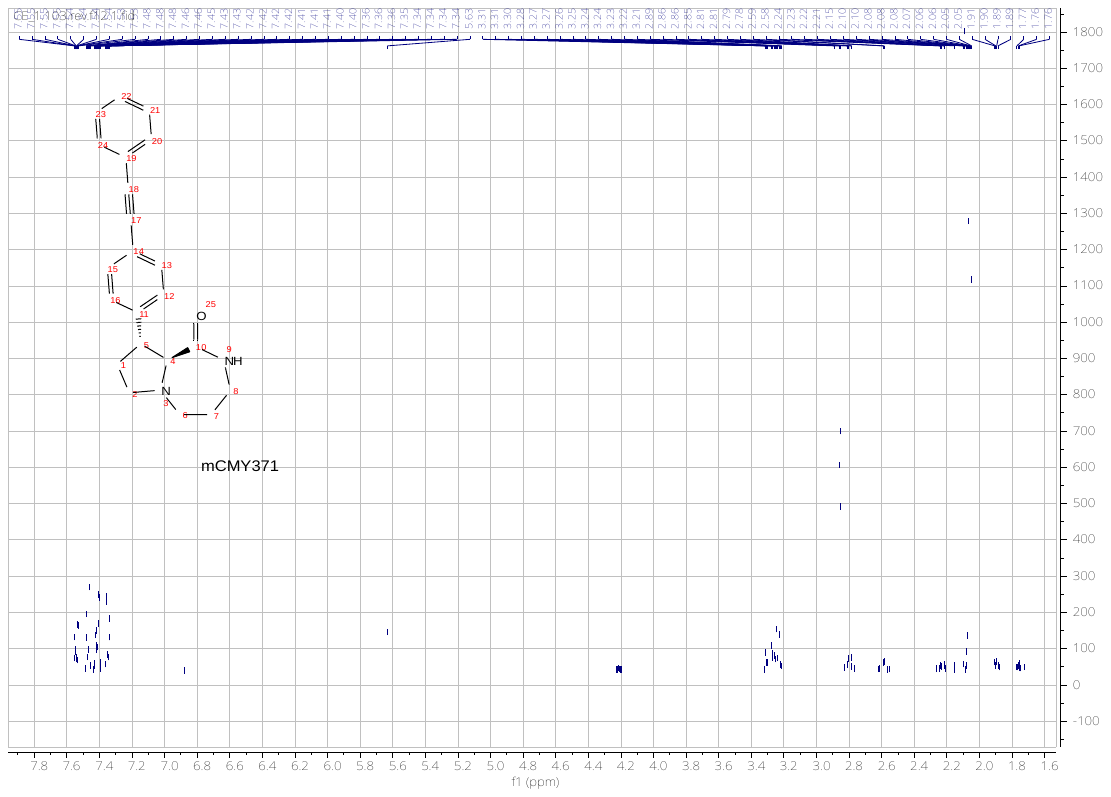

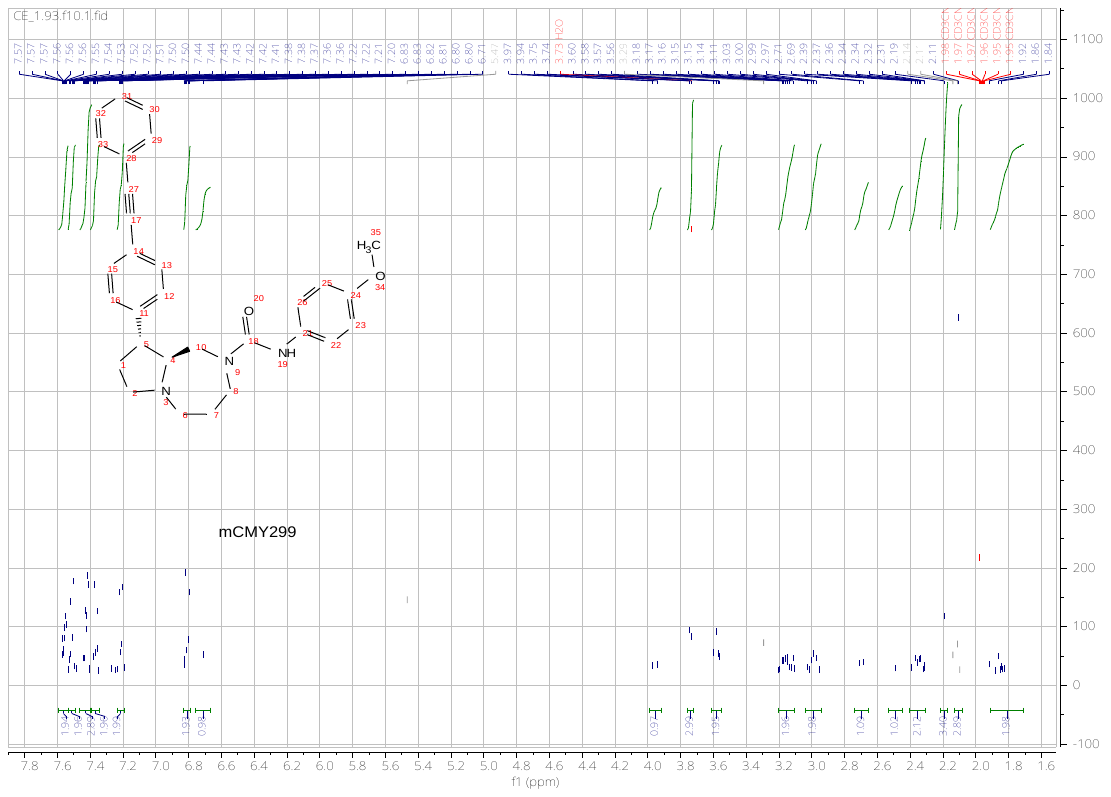

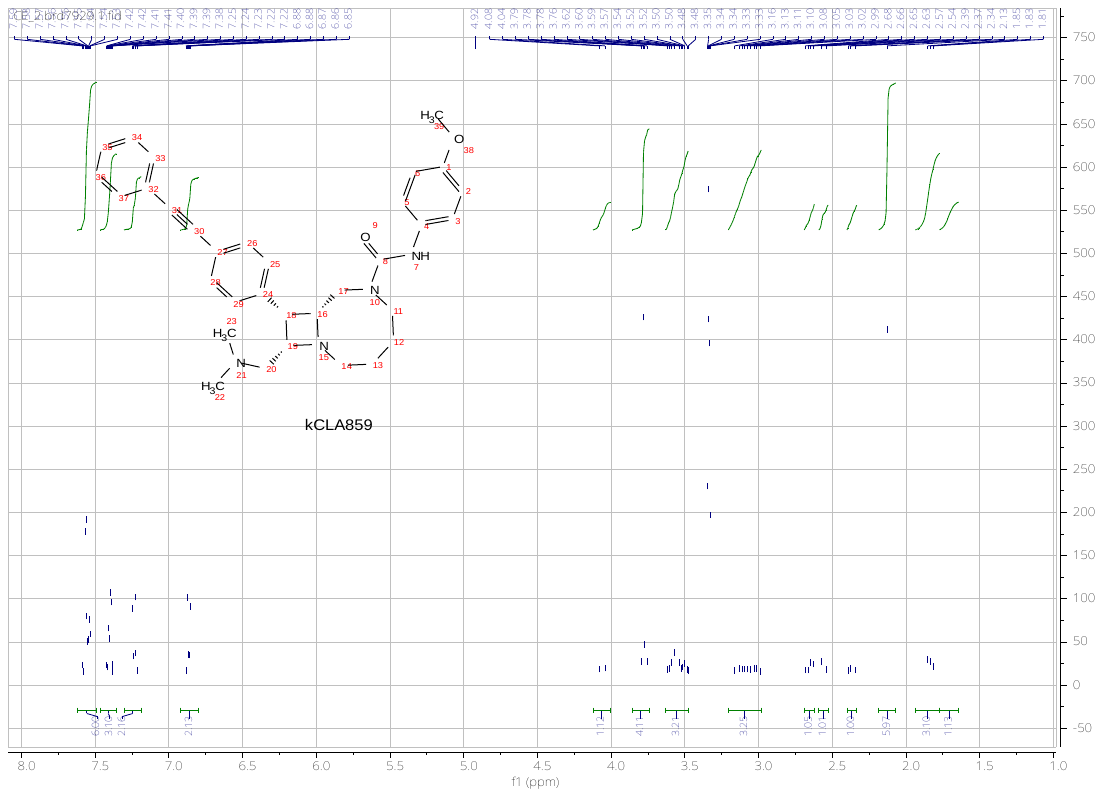

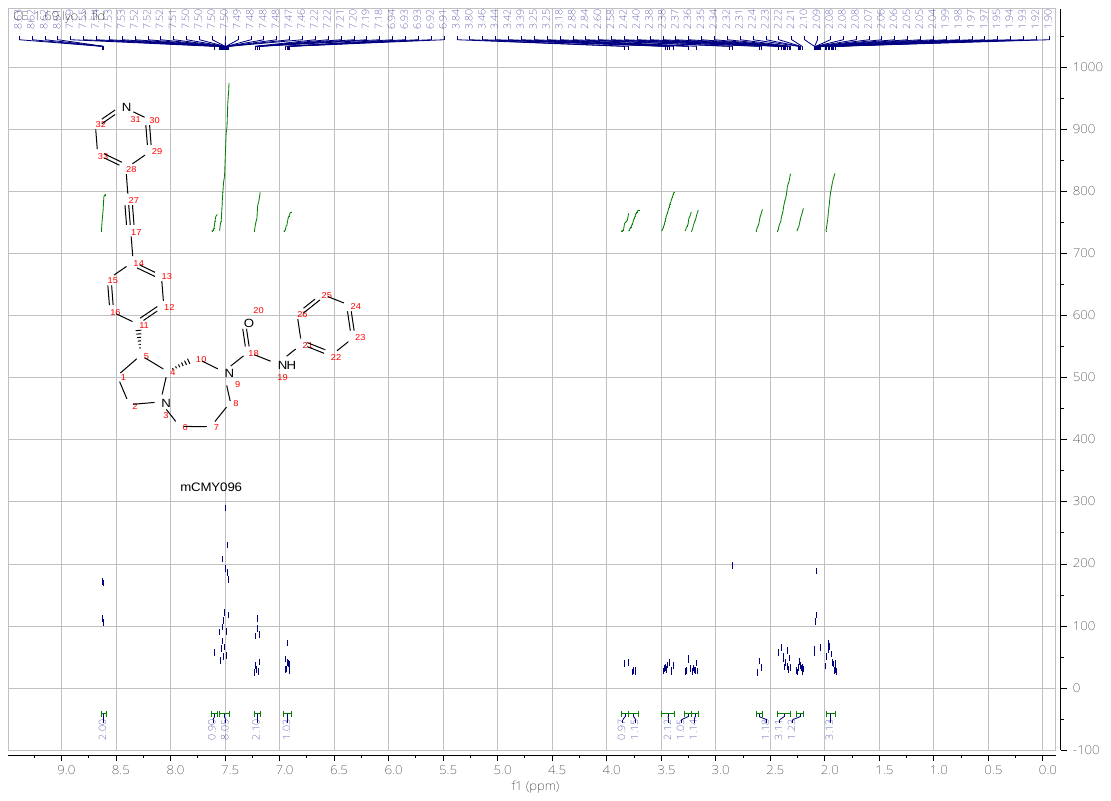

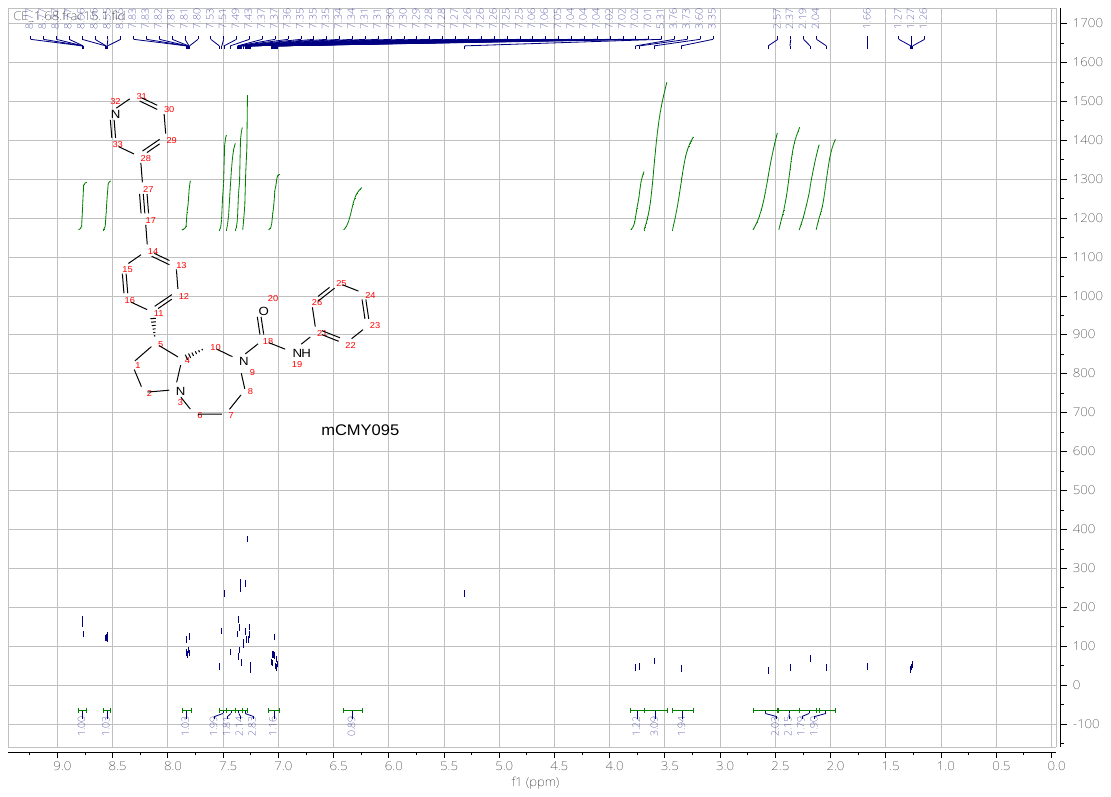

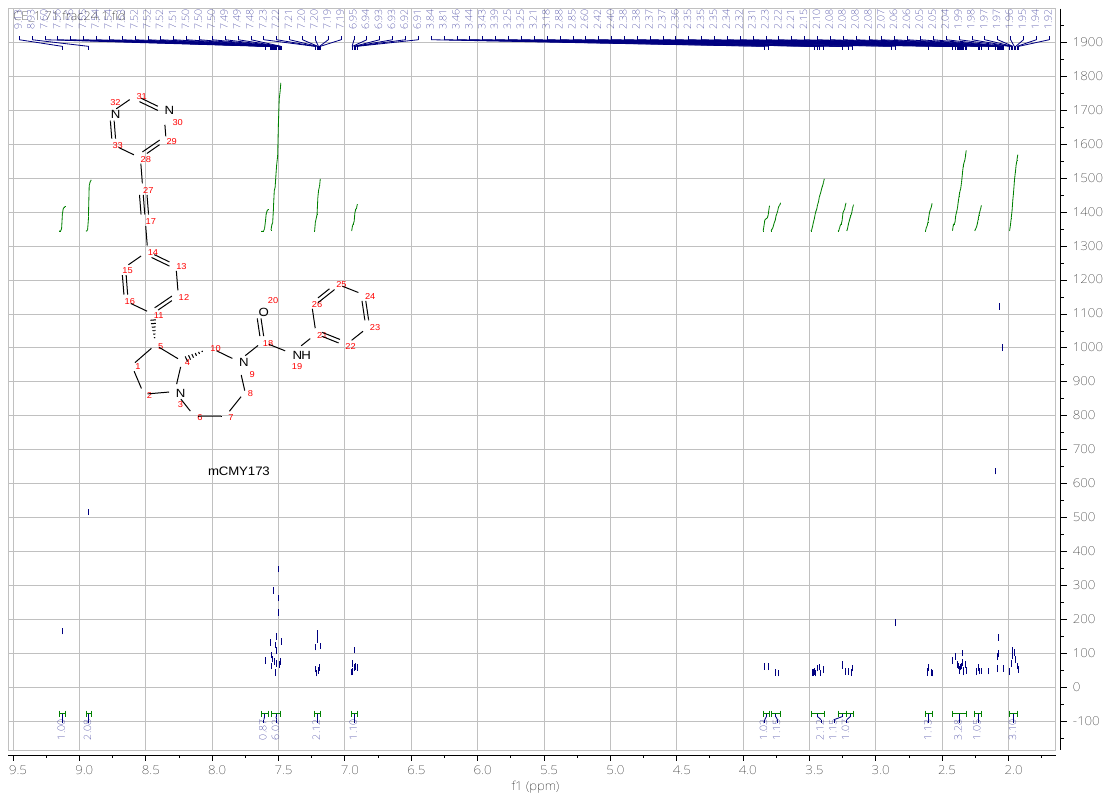

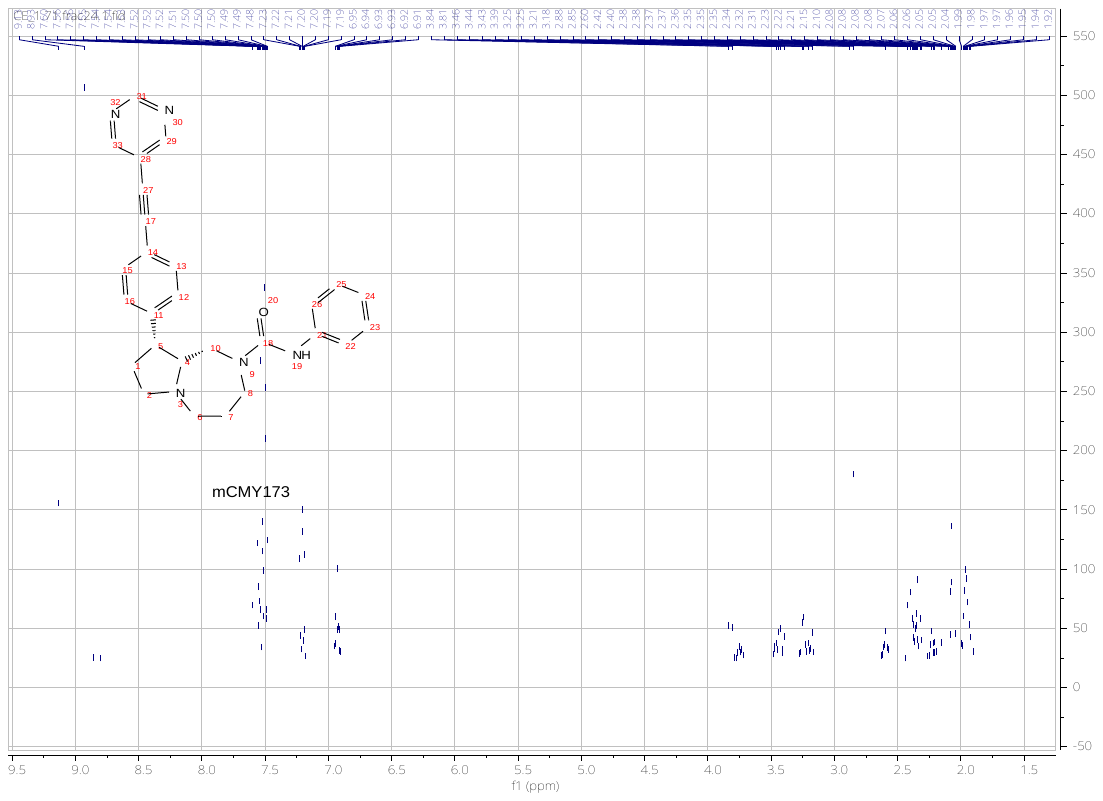

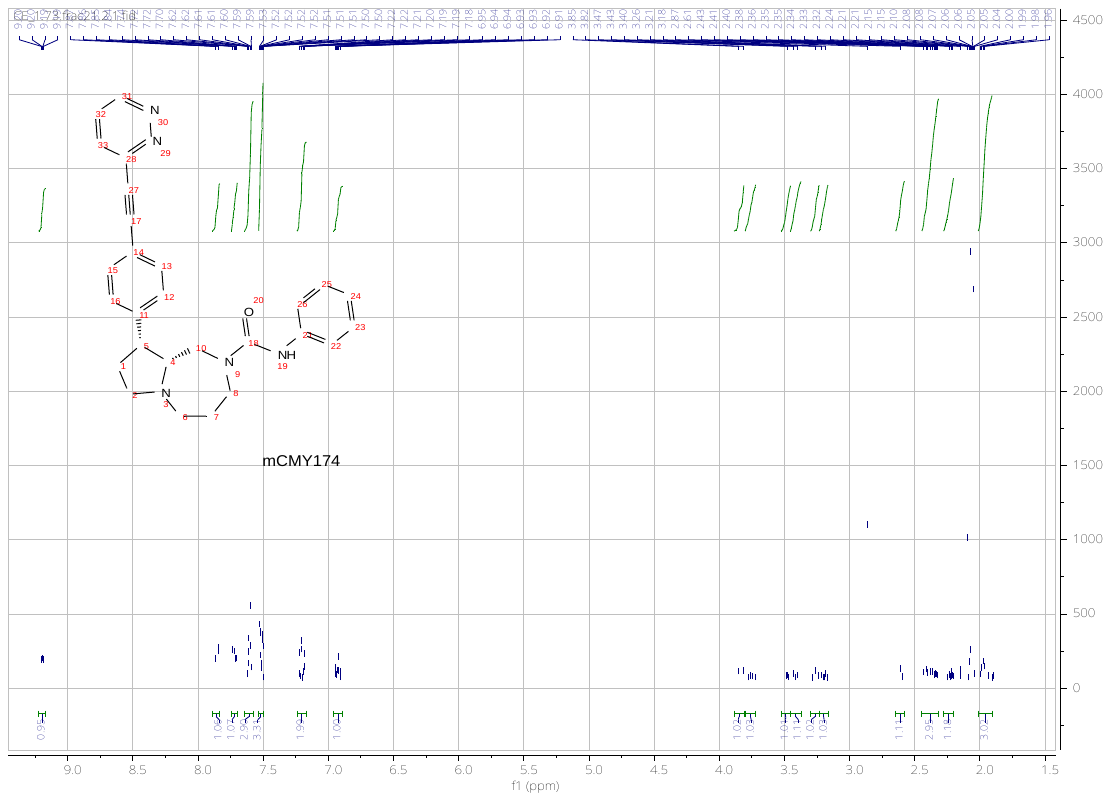

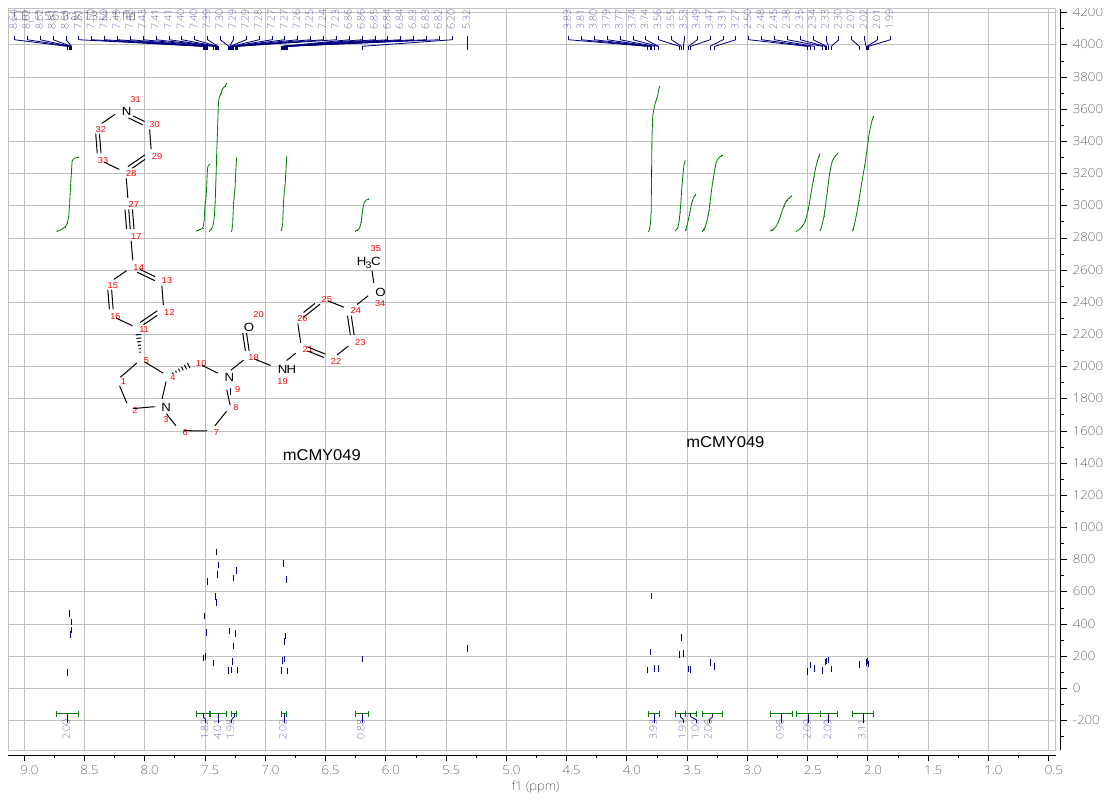

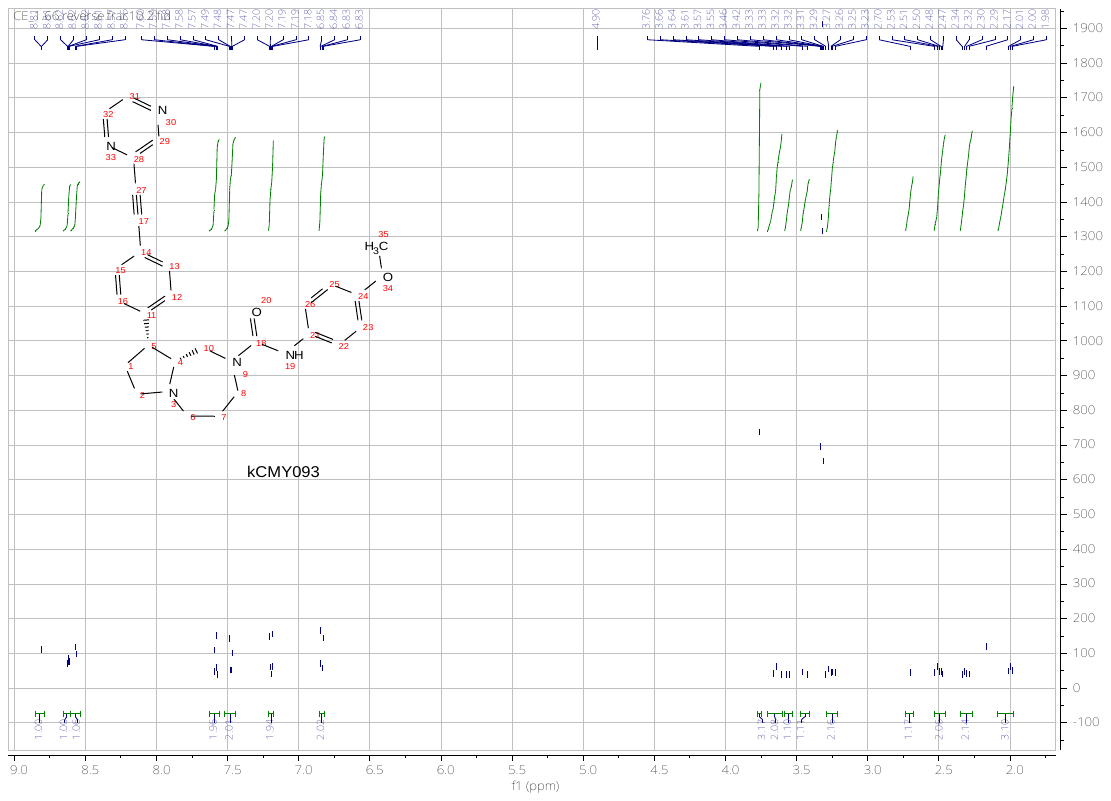

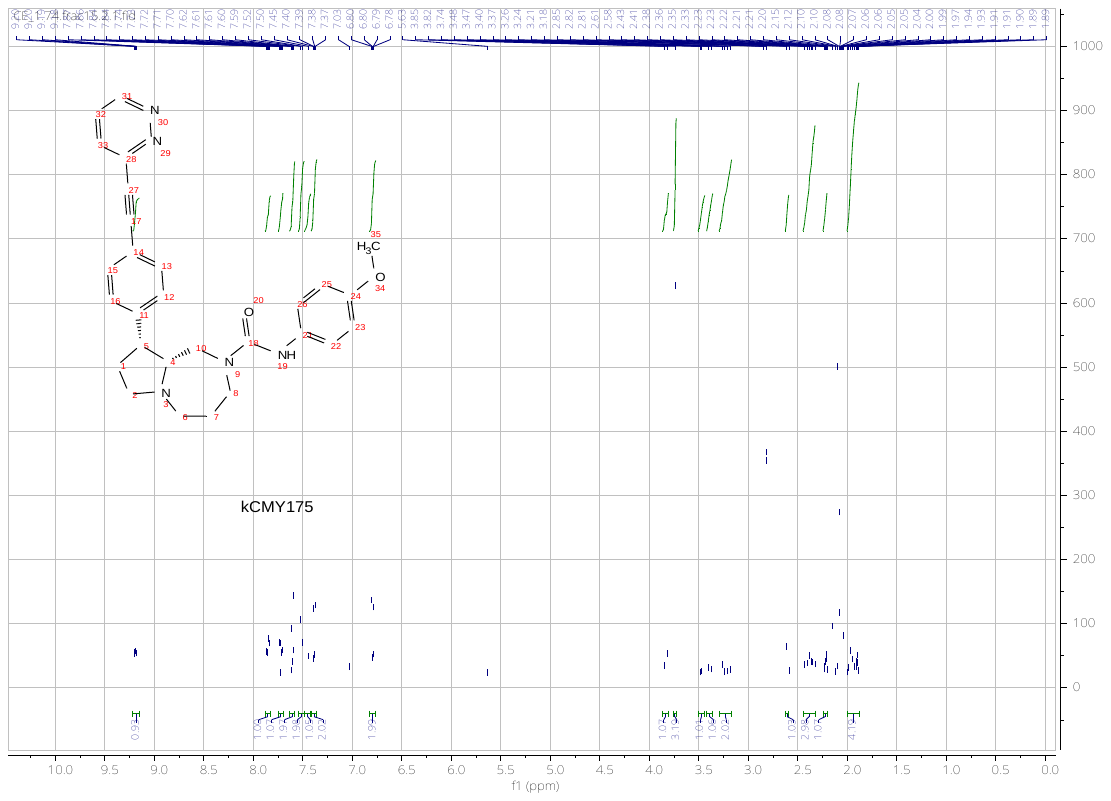

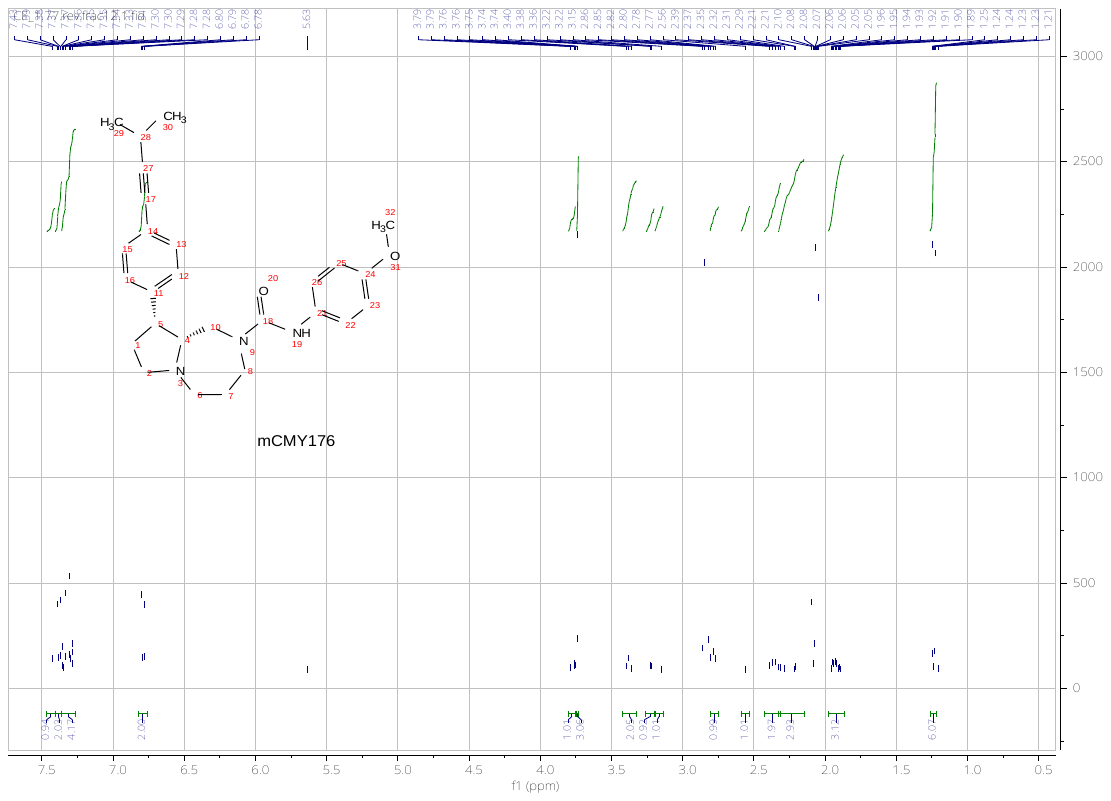

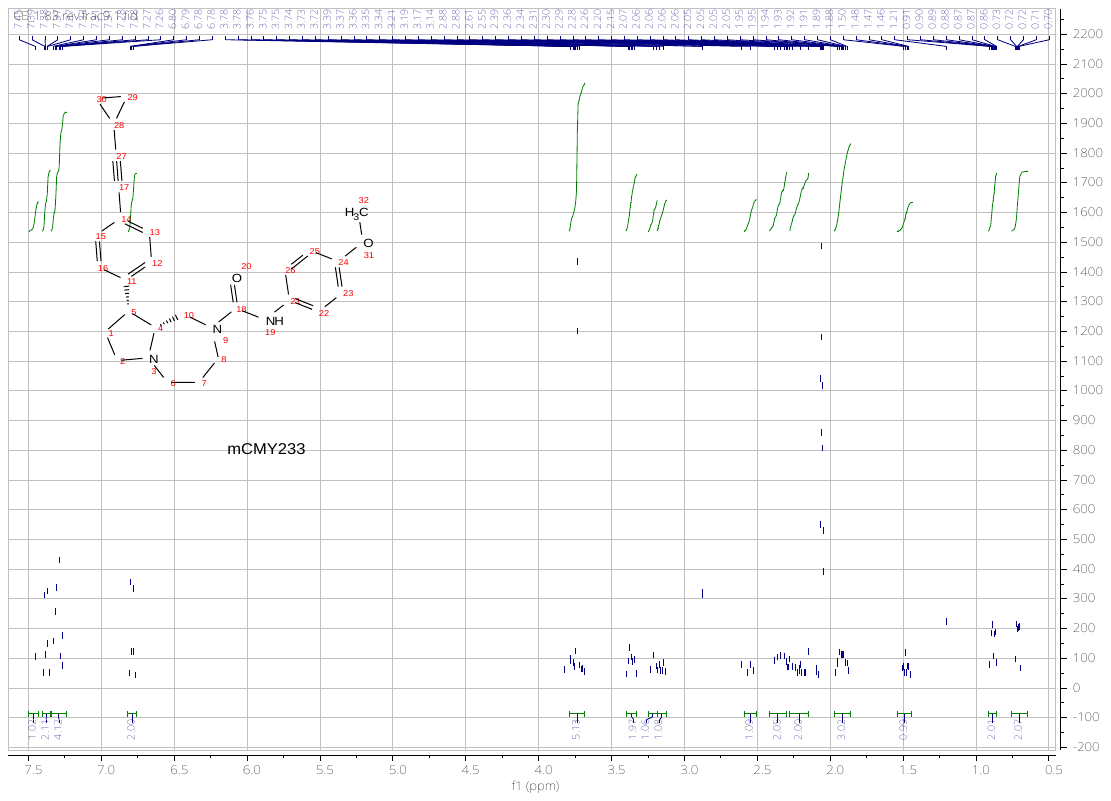

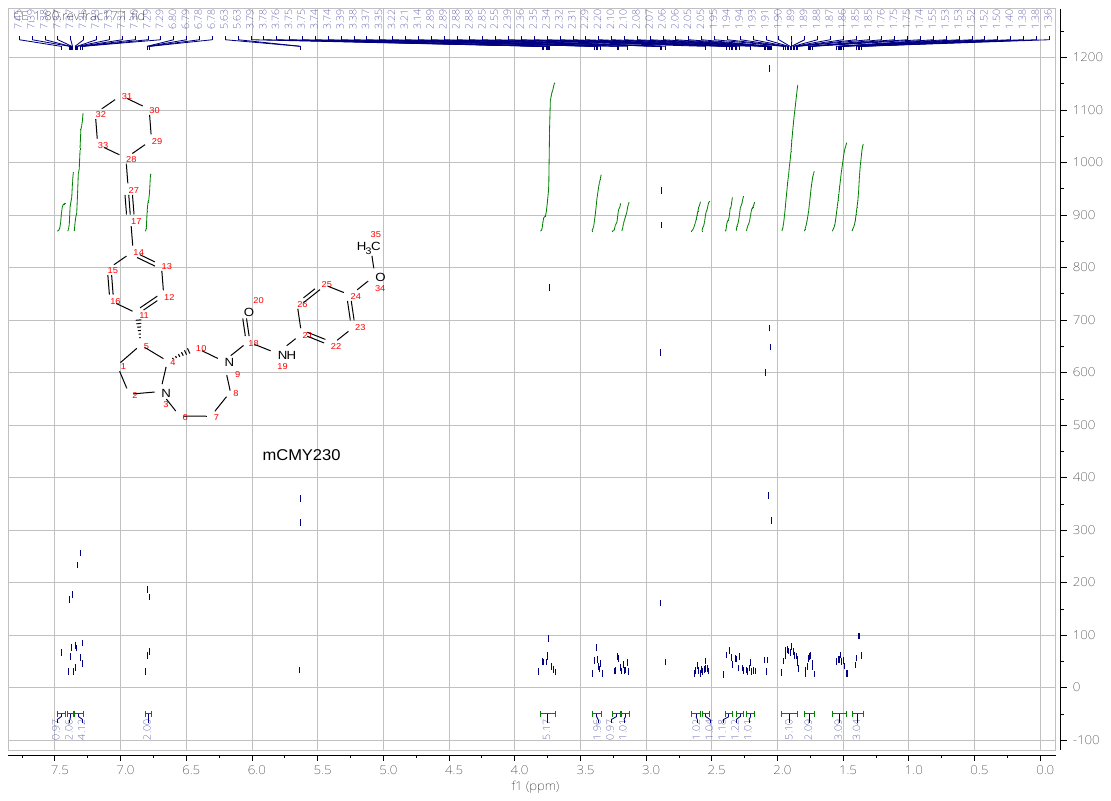

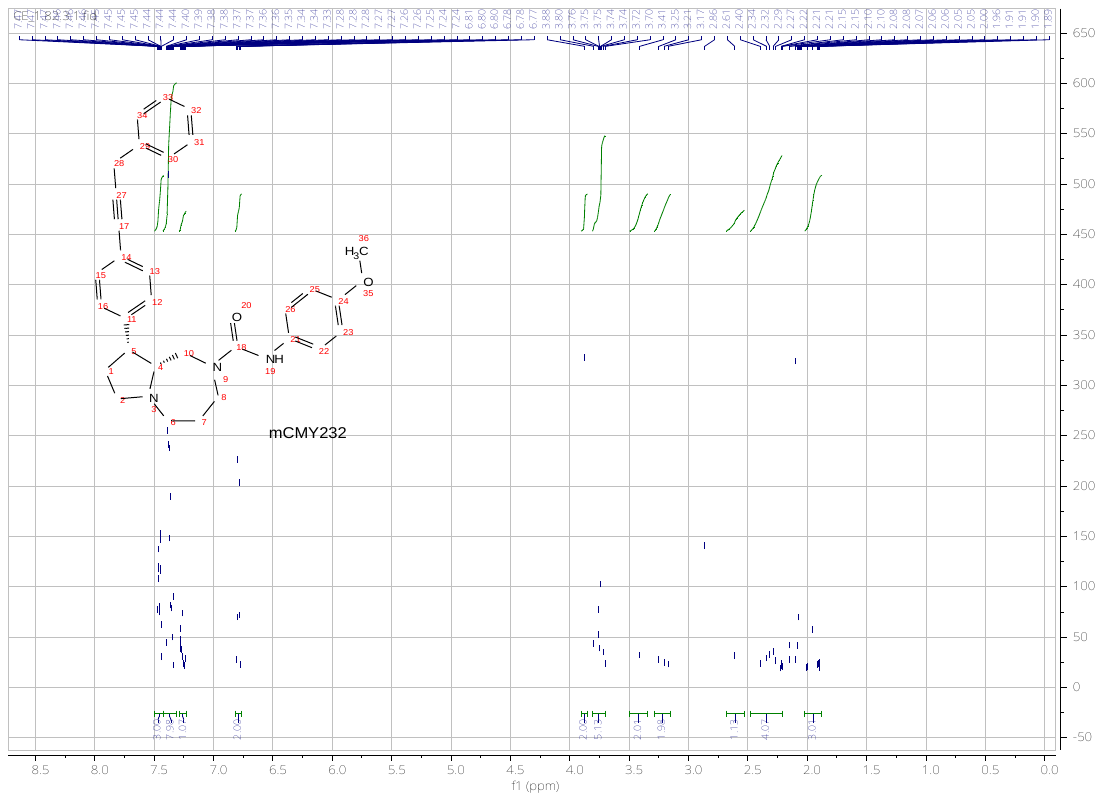

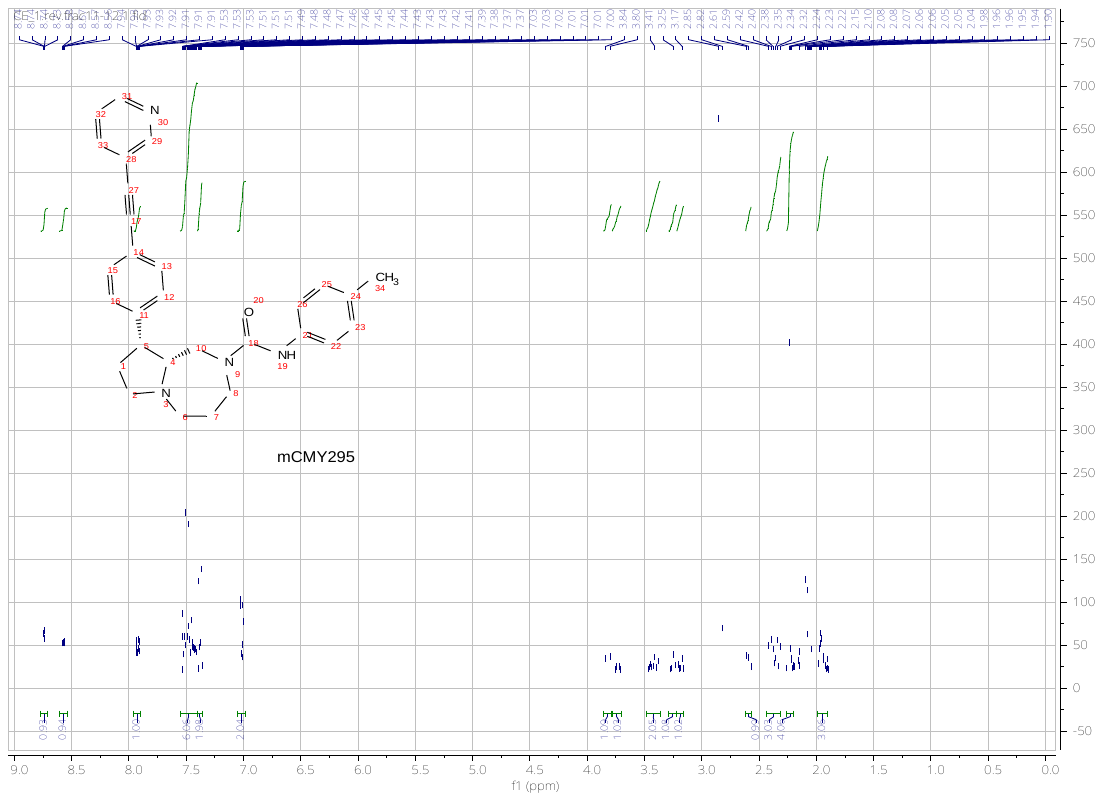

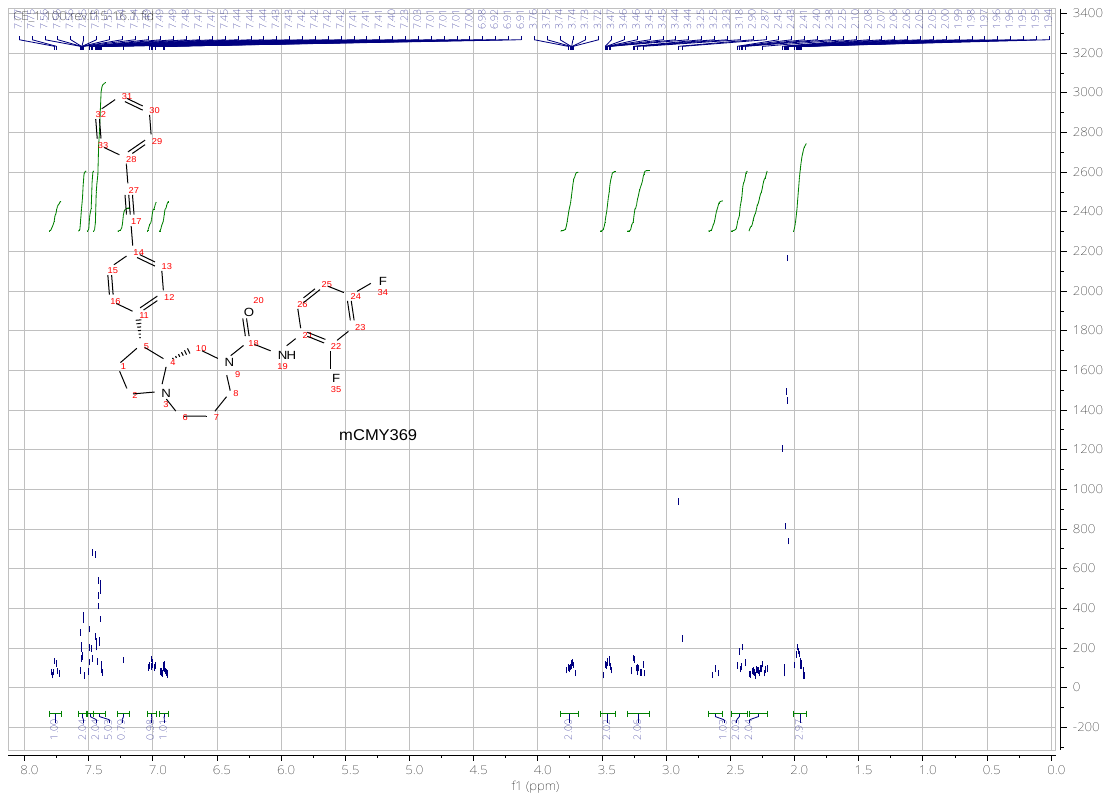

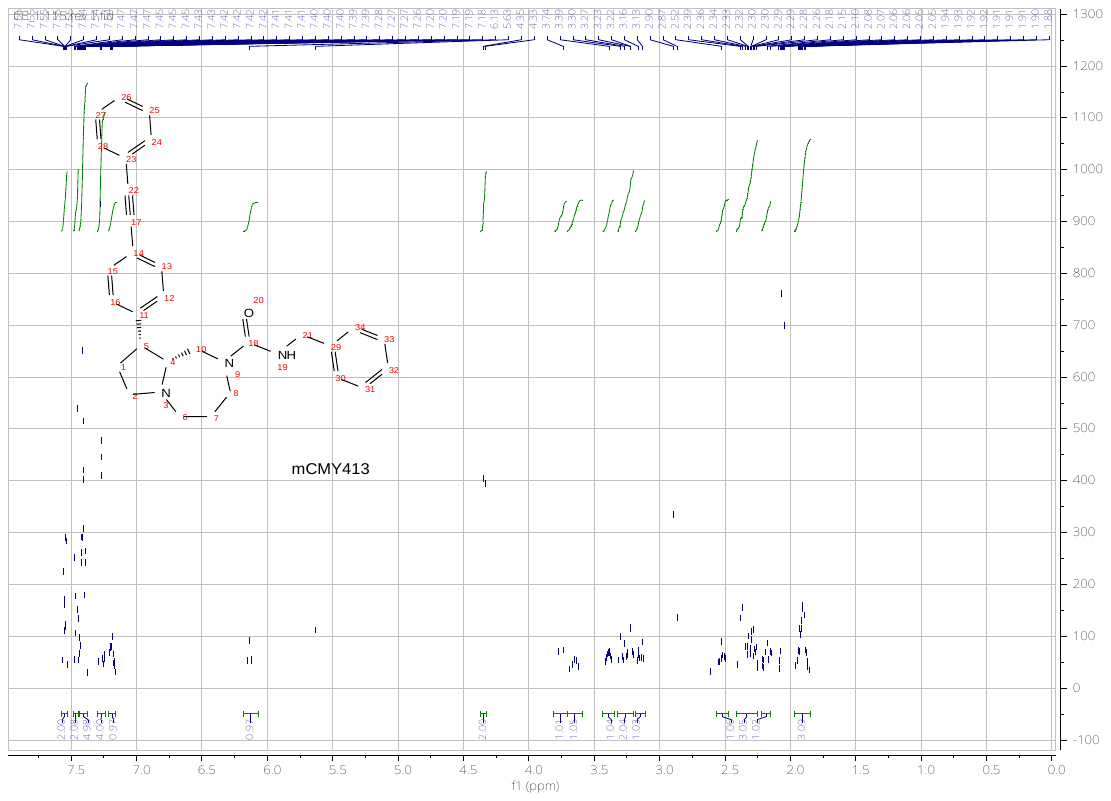

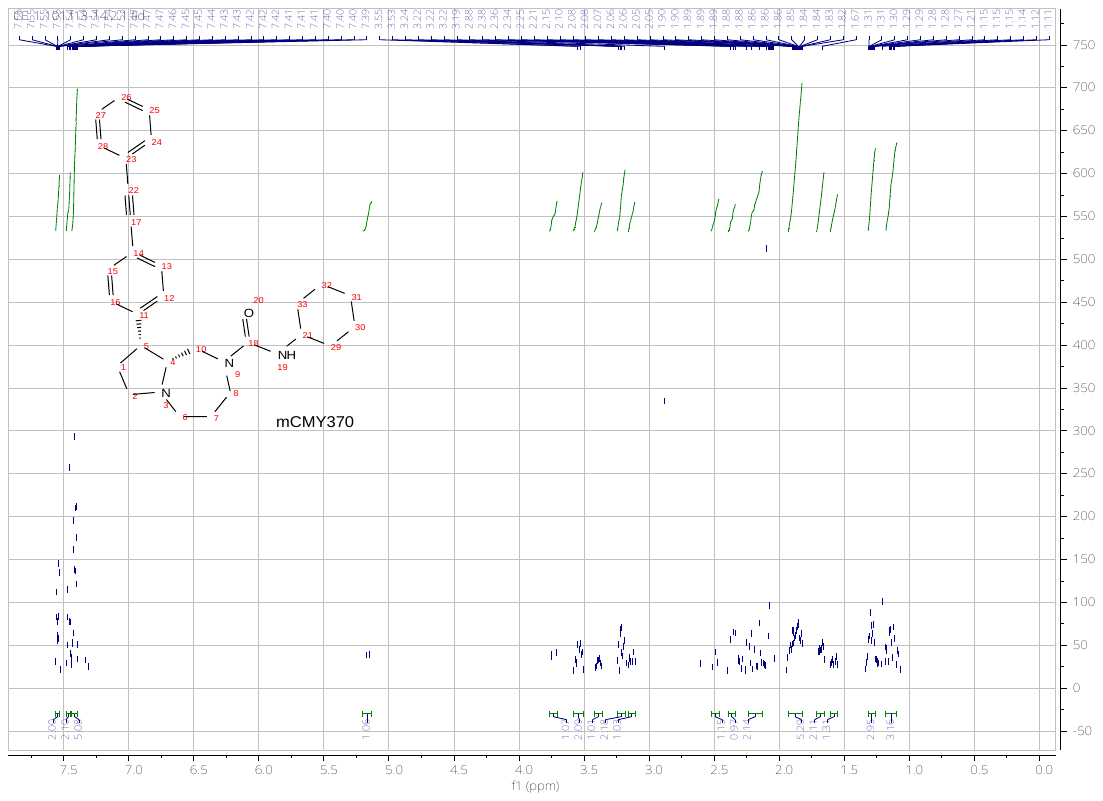

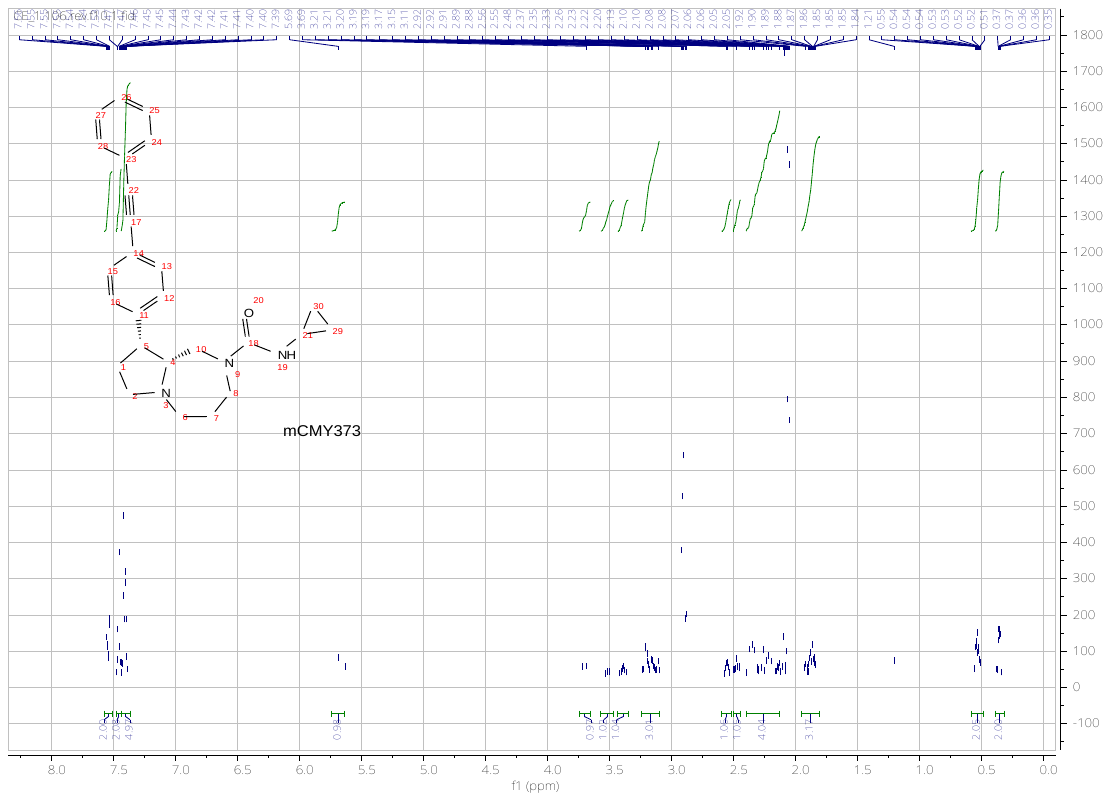

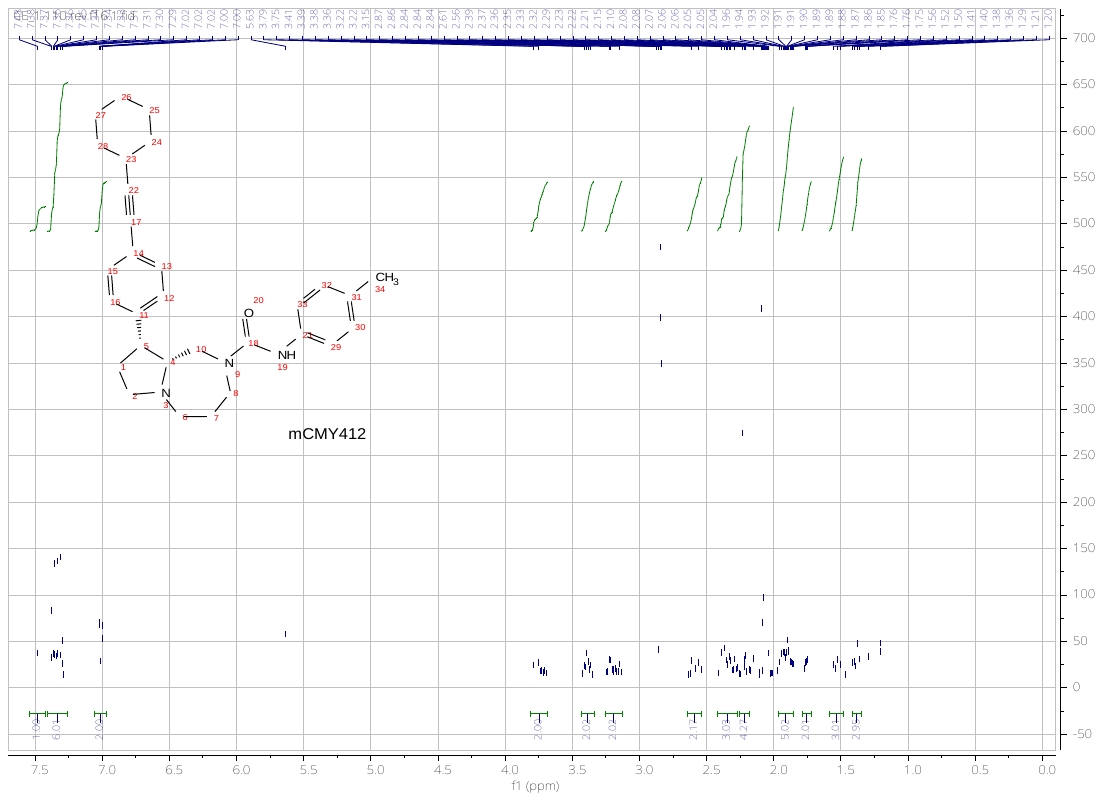

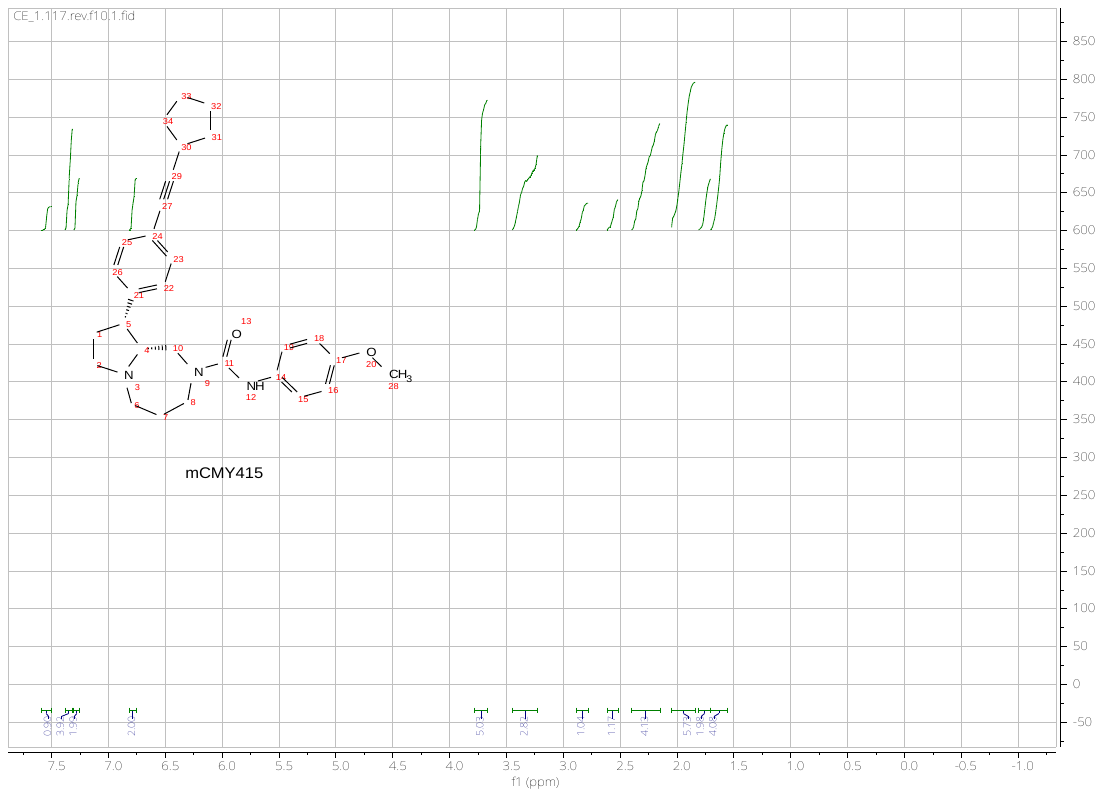

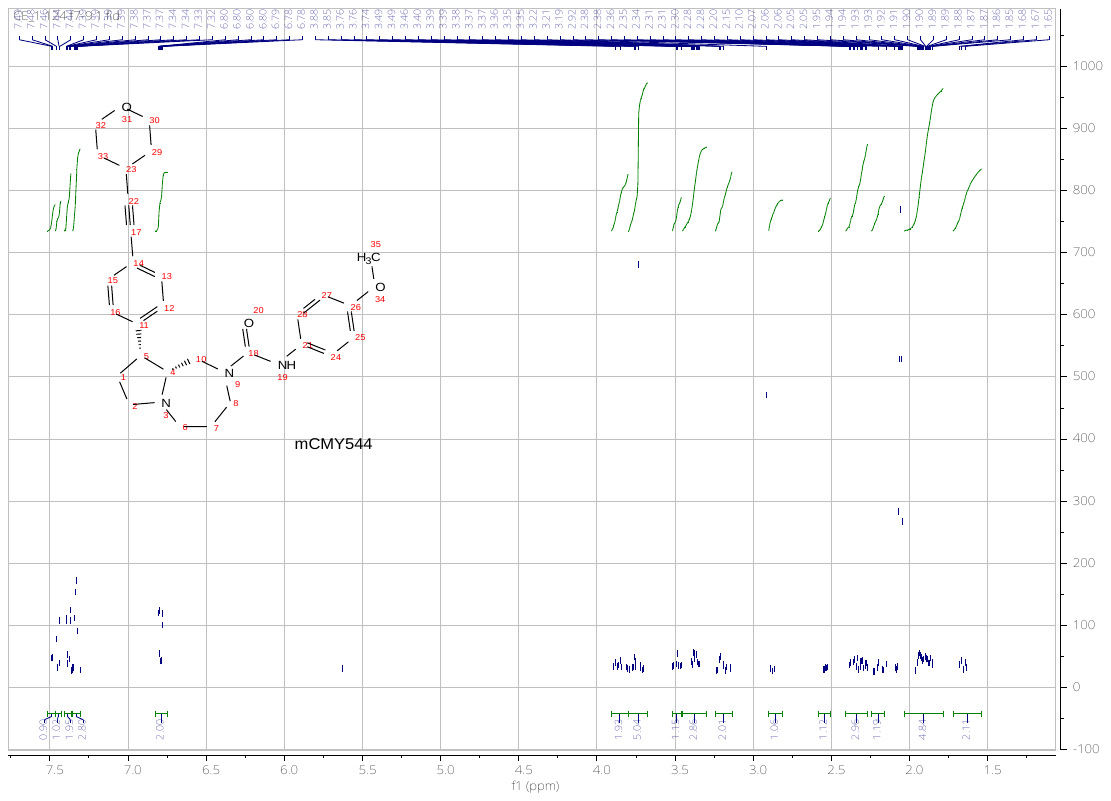

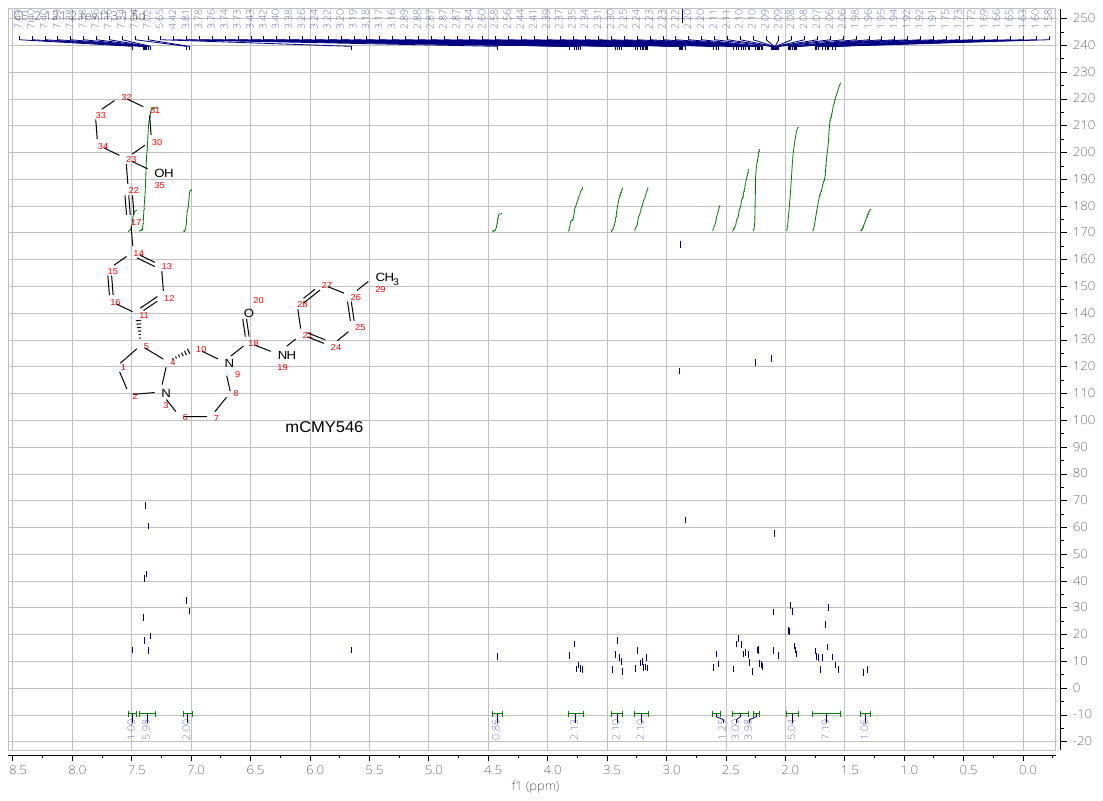

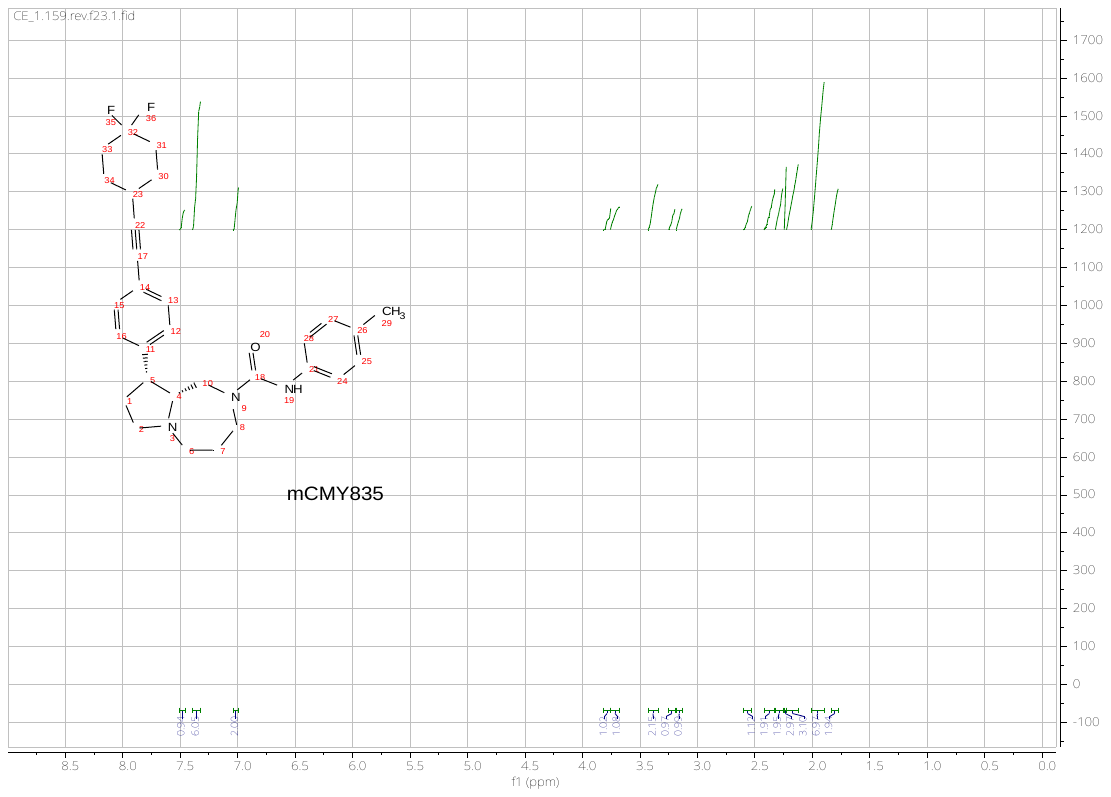

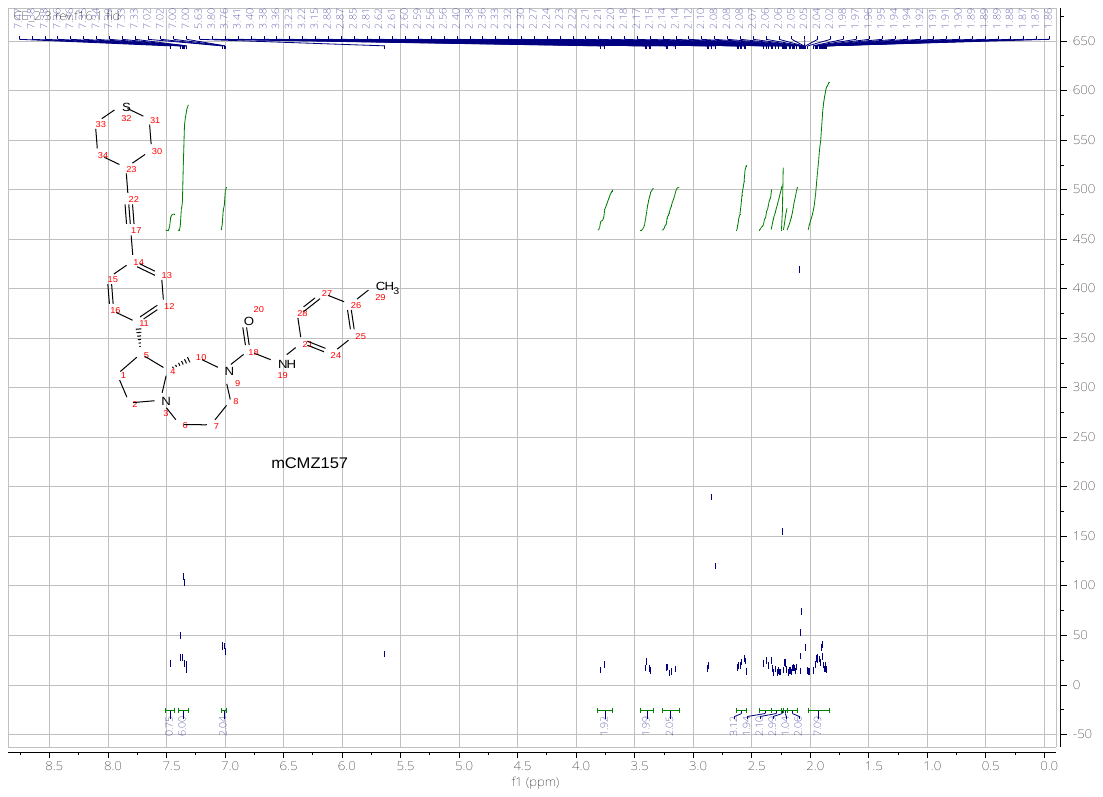

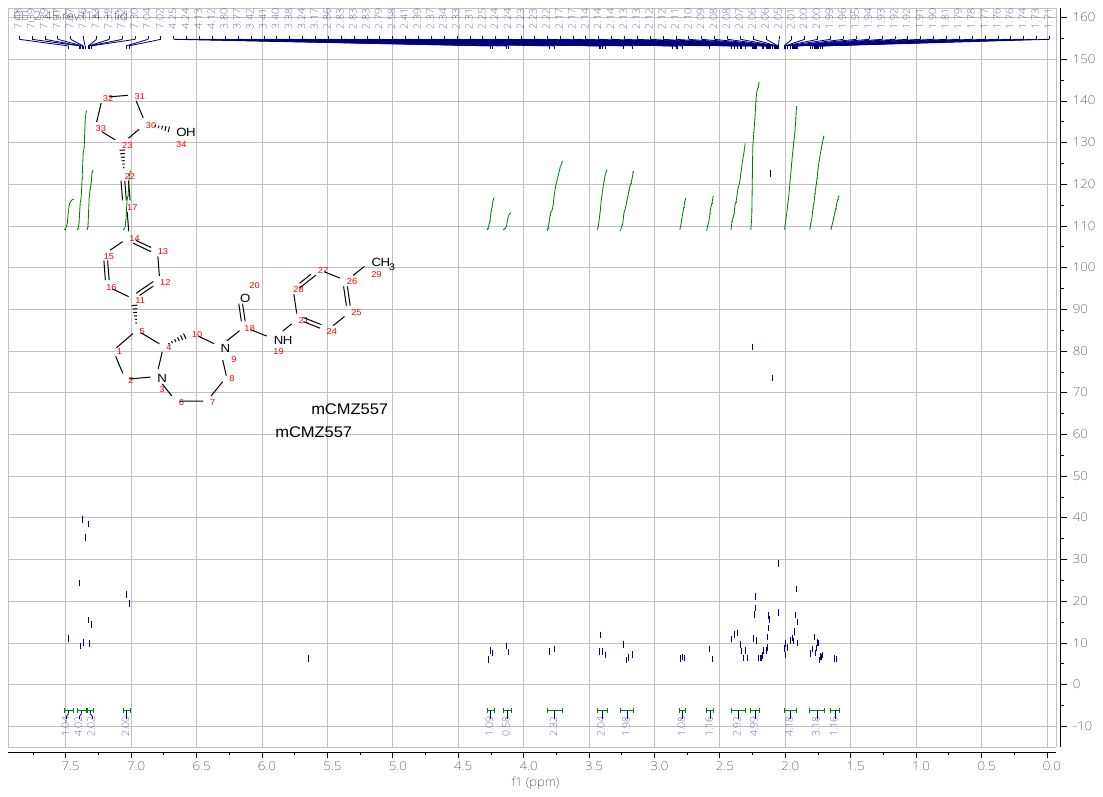

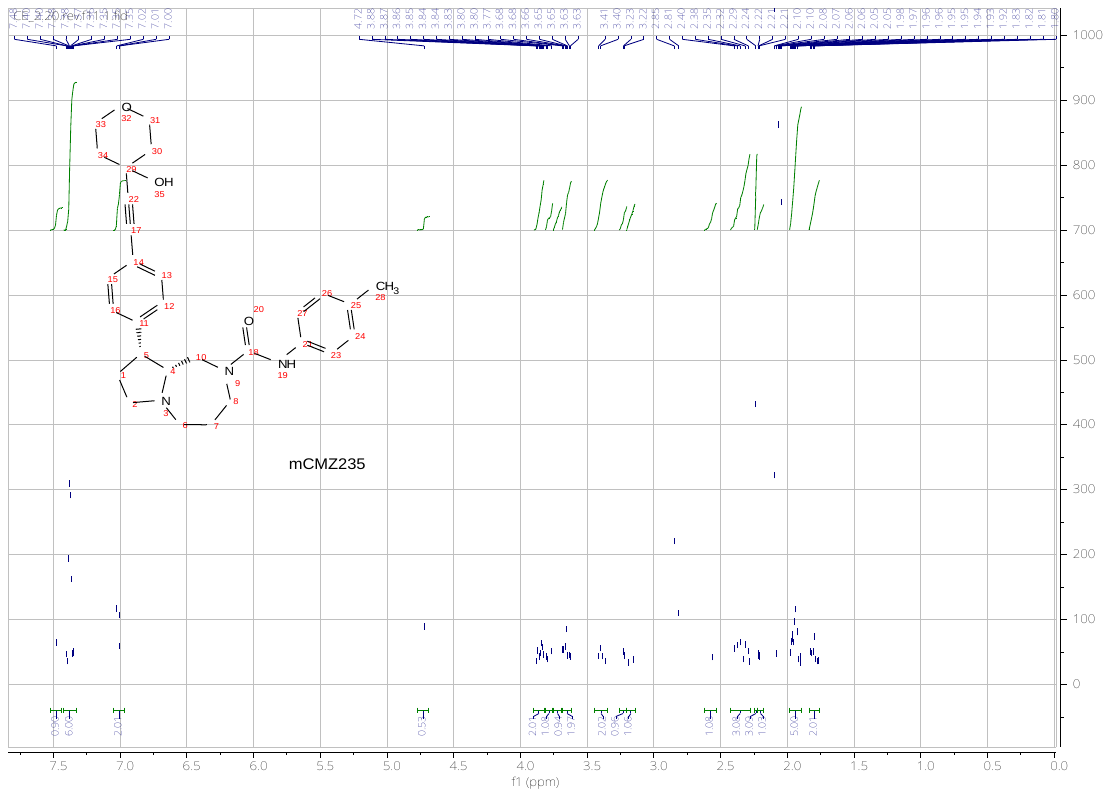
